# Dissociable Roles of Pallidal Neuron Subtypes in Regulating Motor Patterns

**DOI:** 10.1101/2020.08.23.263053

**Authors:** Qiaoling Cui, Arin Pamukcu, Suraj Cherian, Isaac Y. M. Chang, Brianna L. Berceau, Harry S. Xenias, Mathew H. Higgs, Shivakumar Rajamanickam, Yi Chen, Xixun Du, Yu Zhang, Hayley McMorrow, Zachary A. Abecassis, Simina M. Boca, Nicholas J. Justice, Charles J. Wilson, C. Savio Chan

## Abstract

We have previously established that PV^+^ neurons and Npas1^+^ neurons are distinct neuron classes in the GPe— they have different topographical, electrophysiological, circuit, and functional properties. Aside from Foxp2^+^ neurons, which are a unique subclass within the Npas1^+^ class, we lack driver lines that effectively capture other GPe neuron subclasses. In this study, we examined the utility of Kcng4-Cre, Npr3-Cre, and Npy2r-Cre mouse lines (both males and females) for the delineation of GPe neuron subtypes. By using these novel driver lines, we have provided the most exhaustive investigation of electrophysiological studies of GPe neuron subtypes to date. Corroborating our prior studies, GPe neurons can be divided into two statistically distinct clusters that map onto PV^+^ and Npas1^+^ classes. By combining optogenetics and machine learning-based tracking, we showed that optogenetic perturbation of GPe neuron subtypes generated unique behavioral structures. Our findings further highlighted the dissociable roles of GPe neurons in regulating movement and anxiety-like behavior. We concluded that Npr3^+^ neurons and Kcng4^+^ neurons are distinct subclasses of Npas1^+^ neurons and PV^+^ neurons, respectively. Finally, by examining local collateral connectivity, we inferred the circuit mechanisms involved in the motor patterns observed with optogenetic perturbations. In summary, by identifying mouse lines that allow for manipulations of GPe neuron subtypes, we created new opportunities for interrogations of cellular and circuit substrates that can be important for motor function and dysfunction.

**Significance statement:** Within the basal ganglia, the external globus pallidus (GPe) has long been recognized for its involvement in motor control. However, we lacked an understanding of precisely how movement is controlled at the GPe level as a result of its cellular complexity. In this study, by using transgenic and cell-specific approaches, we showed that genetically-defined GPe neuron subtypes have distinct roles in regulating motor patterns. In addition, the *in vivo* contributions of these neuron subtypes are in part shaped by the local, inhibitory connections within the GPe. In sum, we have established the foundation for future investigations of motor function and disease pathophysiology.

## Introduction

The basal ganglia are a network of subcortical structures that are involved in motor control and adaptive behavior. Dysfunction within this circuit can be devastating, as seen in patients afflicted with Parkinson’s disease (PD) (Albin et al., 1989; DeLong, 1990; Graybiel et al., 1994; Mink, 1996; DeLong and Wichmann, 2007; Redgrave et al., 2010; Nelson and Kreitzer, 2014; Jahanshahi et al., 2015; Dudman and Krakauer, 2016; Grillner and Robertson, 2016; Klaus et al., 2019; Park et al., 2020). The external globus pallidus (GPe) is reciprocally connected with the dorsal striatum and subthalamic nucleus (STN) and is known to regulate motor output (Smith et al., 1998; Kita, 2007; Hernandez et al., 2015; Glajch et al., 2016; Hegeman et al., 2016; Pamukcu et al., 2020). Consistent with this idea, decorrelated, phasic changes in GPe neuron activity are observed with normal movements (Anderson and Horak, 1985; Shi et al., 2004; Turner and Anderson, 2005; Jin et al., 2014; Dodson et al., 2015; Mallet et al., 2016). Alterations in the firing pattern of these neurons are associated with hypokinetic motor symptoms in both animal models of PD and human patients (Filion et al., 1991; Hutchison et al., 1994; Nini et al., 1995; Rothblat and Schneider, 1995; Boraud et al., 1998; Raz et al., 2000; Magill et al., 2001; Mallet et al., 2008; Chan et al., 2011; Jaeger and Kita, 2011).

Although the importance of the GPe has long been recognized, our understanding of its cell types, their organization, and their functional properties is limited. While prior studies argue GPe neurons are involved in movement control (Dodson et al., 2015; Zimnik et al., 2015; Glajch et al., 2016; Mastro et al., 2017; Aristieta et al., 2020; Gu et al., 2020; Pamukcu et al., 2020), precisely how they are involved in motor function and dysfunction remains poorly defined. A major hurdle has been imposed by the complexity of the cellular makeup of the GPe. Our past work characterized two principal classes of GPe neurons, parvalbumin-expressing (PV^+^) neurons and Npas1-expressing (Npas1^+^) neurons that account for roughly 50% and 30% of the GPe, respectively (Hernandez et al., 2015; Hegeman et al., 2016; Abecassis et al., 2020). Aside from Foxp2-expressing (Foxp2^+^) neurons that form a unique subclass within the Npas1^+^ class, we lack driver lines that selectively capture other GPe neuron subclasses. The advent of transcriptomic data provided key insights for genetic dissection of GPe neuron subtypes. In particular, pre-existing single-cell transcriptomic studies (Saunders et al., 2018) and *in situ* hybridization data (Lein et al., 2007) suggest the existence of molecular markers that can be utilized but have not been previously explored for the interrogation of GPe neuron subtypes. In this study, we examined the utility of Kcng4-Cre, Npr3-Cre, and Npy2r-Cre mouse lines. Using complementary approaches, such as anatomical, electrophysiological, and behavioral approaches, we concluded that Npr3^+^ neurons and Kcng4^+^ neurons are unique neuron subclasses within the GPe. Critically, capitalizing on newly available transgenic tools, we showed that GPe neuron subtypes were distinct in their roles in regulating motor patterns.

## Methods

### Mice

All procedures were done in accordance with protocols approved by Northwestern University and The University of Texas Health Science Center at Houston Institutional Animal Care and Use Committees and were in compliance with the National Institutes of Health Guide to the Care and Use of Laboratory Animals. Experiments were conducted with the following mouse lines: Adora2a-Cre BAC (A2a-Cre, MMRRC 031168), C57BL/6J (C57, Jax 000664), Drd1a-Cre BAC (D1-Cre, MMRRC 029178), Foxp2-ires-Cre (Foxp2-Cre, Jax 030541), FSF(Frt-STOP-Frt)-LSL-tdTomato (Ai65, Jax 021875), Kcng4-Cre (Jax 029414), Lox-STOP-Lox(LSL)-tdTomato (Ai14, Jax 007914), Npr3-ires2-Cre (Npr3-Cre, Jax 031333), Npy2r-ires-Cre (Npy2r-Cre, Jax 029285), Npas1-Cre-tdTomato BAC (Npas1-Cre, Jax 027718), PV-ires-Cre (PV-Cre, Jax 017320), PV-2A-Flp (PV-Flp, Jax 022730), PV-tdTomato BAC (PV-tdTomato, Jax 027395). To genetically label different neuron populations for histological and electrophysiological studies, the following crosses were made: Foxp2-Cre; PV-tdTomato, Npr3-Cre; Ai14 (Npr3-L-tdTomato), Npy2r-Cre; Ai14 (Npy2r-L-tdTomato), PV-Flp; Npy2r-Cre; Ai65 (PV-Npy2r-FL-tdTomato), PV-Cre; Ai14 (PV-L-tdTomato), PV-tdTomato; Lhx6-eGFP, Npas1-Cre; Lhx6-eGFP, and Npas1-Cre; PV-tdTomato. Only heterozygous and hemizygous mice were used throughout the study to minimize the potential alteration of the phenotypes in mice carrying the transgene alleles (Chan et al., 2012). All mouse lines were maintained by backcrossing with C57BL/6J stock. Mice were group-housed in a 12 h light-dark cycle. Food and water were provided *ad libitum*. The genotypes of all transgenic mice were determined by tail biopsy followed by PCR to identify the presence of the relevant transgenes. Both male and female mice were used in this study.

### Immunolabeling and biocytin visualization

Mice ranging in age from postnatal day 55–80 were anesthetized deeply with a ketamine-xylazine mixture and perfused transcardially first with phosphate-buffered saline (PBS) followed by a fixative containing 4% (w/v) paraformaldehyde (PFA, pH 7.4). Tissue was then postfixed in the same fixative for 2 h at 4 °C. Tissue blocks containing the GPe were sectioned using a vibrating microtome (Leica Instruments) at a thickness of 60 μm. Floating sections were blocked with 10% (v/v) normal goat or donkey serum (Thermo Fisher Scientific) and 0.2% (v/v) Triton X-100 in PBS for 30–60 min, and were subsequently incubated with primary antibodies in the same solution for 16–24 h at 4 °C. Details of the primary antibodies used in this study are listed in **Table 1**. After washes in PBS, the sections were incubated with Alexa-conjugated secondary antibodies (Thermo Fisher Scientific, 1:500 dilution) at room temperature for 2 h. The sections were then washed, mounted with ProLong Antifade mounting medium (Thermo Fisher Scientific), and coverslipped. In a subset of the experiments, DAPI was used to delineate cytoarchitecture of different brain structures. Fluorescent images of injection sites were captured on an epifluorescence microscope (Keyence Corporation) using a 2× or 10× 0.45 numerical aperture (NA) objective. Immunoreactivity in neurons was examined on a laser-scanning confocal microscope (Olympus). For cell quantification, images of the entire GPe were acquired on a laser-scanning confocal microscope with a 60× 1.35 NA oil-immersion objective. Images encompassing the GPe were taken and stitched using FLUOVIEW Viewer (Olympus) or Photoshop (Adobe Systems). Cell counting was performed manually using the cell-counter plugin within Fiji (Schindelin et al., 2012). Cell counts were obtained from optical sections that were captured at 1 µm increments. Neurons were defined by cells that were immuno-positive for HuCD or NeuN (Hernandez et al., 2015; Abecassis et al., 2020). GPe sections from three different equally-spaced (400 µm) mediolateral levels (∼2.5, 2.1, and 1.7 mm from bregma) were sampled and assigned as lateral, intermediate, and medial, respectively (Hernandez et al., 2015; Abecassis et al., 2020). They correspond approximately to sagittal plate 7, 9, and 11 on the Allen reference atlas (http://mouse.brain-map.org/static/atlas). In this study, the GPe is considered to be the structure that spans between the dorsal striatum and the internal capsule, which define the rostral and caudal borders of the GPe on a sagittal plane, respectively. The cytoarchitecture of the ventral border is more ambiguous. For consistency, six non-overlapping z-stacks (212.13 × 212.13 µm) traversing the long axis of the GPe were used to capture its dorsoventral extent.

**Table 1.**
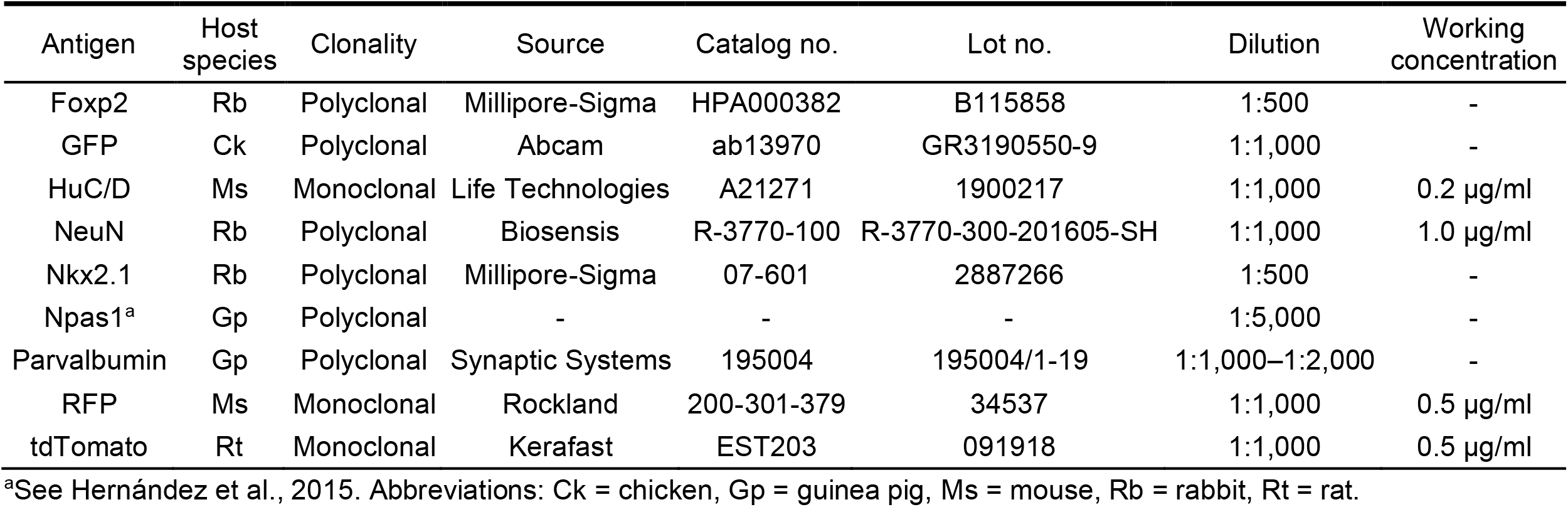
Primary antibodies used in this study.

**Table 2.**
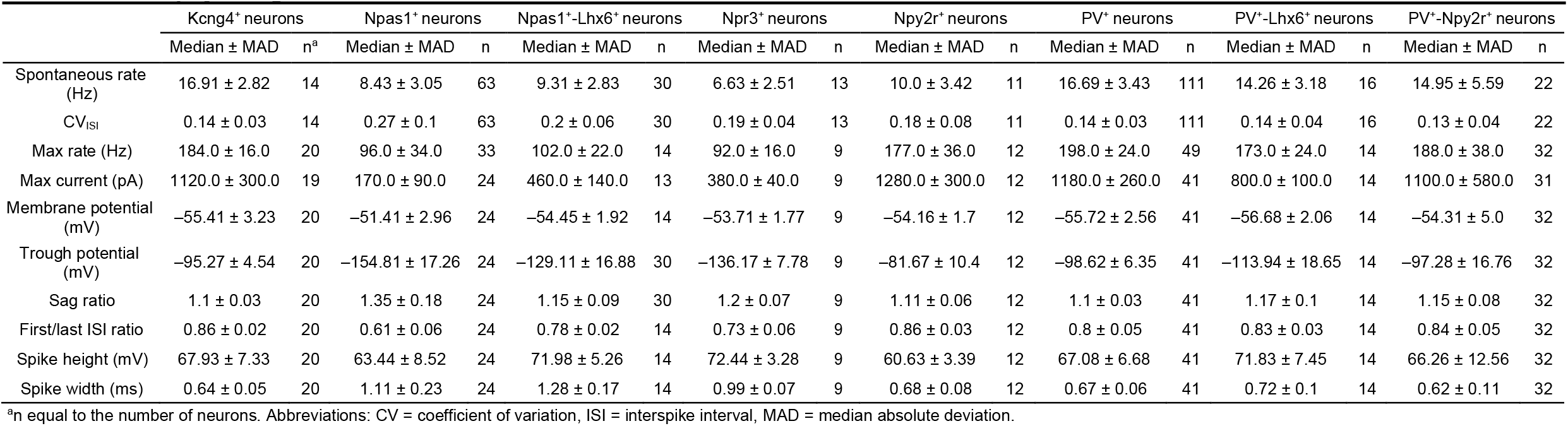
Electrophysiological characteristics of GPe neurons.

|  | Kcng4<sup>+</sup> neurons |  | Npas1<sup>+</sup> neurons |  | Npas1<sup>+</sup>-Lhx6<sup>+</sup> neurons |  | Npr3<sup>+</sup> neurons |  | Npy2r<sup>+</sup> neurons |  | PV<sup>+</sup> neurons |  | PV<sup>+</sup>-Lhx6<sup>+</sup> neurons |  | PV<sup>+</sup>-Npy2r<sup>+</sup> neurons |  |
| --- | --- | --- | --- | --- | --- | --- | --- | --- | --- | --- | --- | --- | --- | --- | --- | --- |
|  | Median ± MAD | n<sup>a</sup> | Median ± MAD | n | Median ± MAD | n | Median ± MAD | n | Median ± MAD | n | Median ± MAD | n | Median ± MAD | n | Median ± MAD | n |
| Spontaneous rate (Hz) | 16.91 ± 2.82 | 14 | 8.43 ± 3.05 | 63 | 9.31 ± 2.83 | 30 | 6.63 ± 2.51 | 13 | 10.0 ± 3.42 | 11 | 16.69 ± 3.43 | 111 | 14.26 ± 3.18 | 16 | 14.95 ± 5.59 | 22 |
| CV<sub>ISI</sub> | 0.14 ± 0.03 | 14 | 0.27 ± 0.1 | 63 | 0.2 ± 0.06 | 30 | 0.19 ± 0.04 | 13 | 0.18 ± 0.08 | 11 | 0.14 ± 0.03 | 111 | 0.14 ± 0.04 | 16 | 0.13 ± 0.04 | 22 |
| Max rate (Hz) | 184.0 ± 16.0 | 20 | 96.0 ± 34.0 | 33 | 102.0 ± 22.0 | 14 | 92.0 ± 16.0 | 9 | 177.0 ± 36.0 | 12 | 198.0 ± 24.0 | 49 | 173.0 ± 24.0 | 14 | 188.0 ± 38.0 | 32 |
| Max current (pA) | 1120.0 ± 300.0 | 19 | 170.0 ± 90.0 | 24 | 460.0 ± 140.0 | 13 | 380.0 ± 40.0 | 9 | 1280.0 ± 300.0 | 12 | 1180.0 ± 260.0 | 41 | 800.0 ± 100.0 | 14 | 1100.0 ± 580.0 | 31 |
| Membrane potential (mV) | -55.41 ± 3.23 | 20 | -51.41 ± 2.96 | 24 | -54.45 ± 1.92 | 14 | -53.71 ± 1.77 | 9 | -54.16 ± 1.7 | 12 | -55.72 ± 2.56 | 41 | -56.68 ± 2.06 | 14 | -54.31 ± 5.0 | 32 |
| Trough potential (mV) | -95.27 ± 4.54 | 20 | -154.81 ± 17.26 | 24 | -129.11 ± 16.88 | 30 | -136.17 ± 7.78 | 9 | -81.67 ± 10.4 | 12 | -98.62 ± 6.35 | 41 | -113.94 ± 18.65 | 14 | -97.28 ± 16.76 | 32 |
| Sag ratio | 1.1 ± 0.03 | 20 | 1.35 ± 0.18 | 24 | 1.15 ± 0.09 | 30 | 1.2 ± 0.07 | 9 | 1.11 ± 0.06 | 12 | 1.1 ± 0.03 | 41 | 1.17 ± 0.1 | 14 | 1.15 ± 0.08 | 32 |
| First/last ISI ratio | 0.86 ± 0.02 | 20 | 0.61 ± 0.06 | 24 | 0.78 ± 0.02 | 14 | 0.73 ± 0.06 | 9 | 0.86 ± 0.03 | 12 | 0.8 ± 0.05 | 41 | 0.83 ± 0.03 | 14 | 0.84 ± 0.05 | 32 |
| Spike height (mV) | 67.93 ± 7.33 | 20 | 63.44 ± 8.52 | 24 | 71.98 ± 5.26 | 14 | 72.44 ± 3.28 | 9 | 60.63 ± 3.39 | 12 | 67.08 ± 6.68 | 41 | 71.83 ± 7.45 | 14 | 66.26 ± 12.56 | 32 |
| Spike width (ms) | 0.64 ± 0.05 | 20 | 1.11 ± 0.23 | 24 | 1.28 ± 0.17 | 14 | 0.99 ± 0.07 | 9 | 0.68 ± 0.08 | 12 | 0.67 ± 0.06 | 41 | 0.72 ± 0.1 | 14 | 0.62 ± 0.11 | 32 |

<sup>a</sup>n equal to the number of neurons. Abbreviations: CV = coefficient of variation, ISI = interspike interval, MAD = median absolute deviation.

To visualize biocytin GPe neurons, 0.2% (w/v) biocytin was included in the internal solution and recorded in the whole-cell mode for at least 30 minutes (see below). Recovered slices were flat-mounted on nitrocellulose filters (Millipore) and fixed overnight in 4% (w/v) paraformaldehyde (PFA, pH 7.4) at 4 °C. After several washes in PBS, slices were reacted three times in freshly prepared 0.5% (w/v) NaBH_4_ in H_2_O. Slices were then incubated in fresh Scale (Hama et al., 2011) solution (4 M urea, 10% (v/v) glycerol and 0.1% Triton X-100) for 10 days at 4 °C. Subsequently, slices were washed in PBST (1% Triton X-100 in PBS) and blocked in 10% (v/v) normal goat serum and PBST for 45 min. After several washes, slices were reacted with 2 μg/ml streptavidin-Alexa Fluor 647 (Thermo Fisher Scientific) in blocking solution overnight at 4 °C in the dark. Slices were washed in PBS followed by Scale solution. Sections were mounted, air-dried, and then coverslipped in ProLong Gold mounting medium (Thermo Fisher Scientific). Serial optical sections (Z-stacks) were acquired on a laser-scanning confocal microscope (Olympus) with a 60× 1.35 NA oil-immersion objective (Olympus) at 1 μm intervals and stitched together using Fiji.

### Stereotaxic injections and fiber implantations

For *in vivo* optogenetic experiments, mice aged postnatal day 28–35 were anesthetized in an isoflurane induction chamber at 3–4% isoflurane and immobilized on a stereotaxic frame (David Kopf Instruments). Anesthesia was maintained using 1–2% isoflurane. The scalp was opened using a scalpel and a small craniotomy (1 mm diameter) was made with a dental drill (Osada). Adeno-associated viruses (AAVs) were infused with calibrated 5 µl glass pipettes (VWR) pulled to have a tip diameter of 3 µm. The injection needle was left *in situ* for 5–10 min following the end of the injection to maximize tissue retention of AAV and decrease capillary spread upon pipette withdrawal. Experiments were performed 4–6 weeks after stereotaxic surgeries. The accuracy of injections was visually inspected under epifluorescence microscopy in *ex vivo* slices or histologically verified *post hoc*.

To allow for optogenetic manipulations, fiber optic cannulae were implanted bilaterally at the region of interest (Pamukcu et al., 2020) three weeks after stereotaxic injections. Each fiber cannula was assembled with an optical fiber (0.66 NA, 250 µm core) (Prizmatix) secured into a zirconia ferrule (1.25 mm O.D.) (Prizmatix) with non-fluorescent epoxy (Thorlabs). To determine the intensity of light exiting fibers, the output of fibers was measured with a power meter (Thorlabs). Cannulae were fixed to the skull using dental cement (Parkell).

### Behavioral testing

*In vivo* optogenetic interrogation was performed 1–2 weeks after fiber implantations. Behavioral tests were performed between 2:00 pm and 7:00 pm. *In vivo* optogenetic experiments were performed in an opaque white plastic open-field box (28 cm × 28 cm), which was cleaned with 70% ethanol. On the day prior to optogenetic experiments, each mouse was allowed to acclimate to the open-field box for twenty minutes, with a fiber optic cable attached to the implant via a rotary joint (Prizmatix). Pre-trial, basal ambulatory activity was collected for five minutes at the beginning of the behavioral session. For ChR2 activation, mice were subjected to either a single, sustained light pulse or a patterned pulse train (5 ms pulses at 20 Hz) delivered for 10 s. For GtACR2 activation, only the sustained protocol was used. A blue (peak, ∼455 nm) LED (Prizmatix) was used to activate both opsins. The light power used for opsin activation was 12–18 mW measured at the fiber tip. Stimuli were delivered with a one-minute intertrial interval. An overhead camera (Logitech) was used to acquire mouse behavior in the open field arena at 30 fps, 640 pixels × 240 pixels per frame. Videos were subsequently cropped and downsampled to 10 fps at full resolution for markerless tracking.

### Behavioral tracking and classification

DeepLabCut (https://github.com/DeepLabCut/) (Mathis et al., 2018; Nath et al., 2019) was used for tracking body parts of mice in an open field arena. Eight body parts including the nose, ears, body center, side laterals (hip-joints), tail base, and tail end were labeled in top-down view videos (**Figure 6c**). To create the training dataset, 1,674 distinct frames from 50 video recordings of open-field behavior were manually annotated. We used MobileNetV2-1-based network (Sandler et al., 2018; Mathis et al., 2019) with default parameters. The network was trained and refined for five rounds using default multi-step learning rates. Each round consists of 240,000–1,000,000 iterations, and the default multi-step learning rates were used. This trained network has a test error of 1.13 pixels and a training error of 4.82 pixels. Predictions of X-Y coordinates were processed using a median filter with a rolling window of five frames before further analysis. To cross-validate our network performance, speeds computed from DeepLabCut and Ethovision data were compared; near-identical results were obtained (see Results). This network was then used to analyze all videos in this study.

To categorize motor behavior, DeepLabCut tracking data were first calibrated; the pixel-to-cm conversion for each video was determined by comparing the width of the arena in pixels to the actual width of the arena (28 cm). Based on the calibrated X-Y coordinates of labeled body parts, a set of movement metrics was generated for each frame. Mouse speed was measured as the body center speed. Body angle was defined as tail base–nose angle relative to the body center. An angle < 180° indicates the body axis is skewed to the right, and an angle > 180° indicates the body axis is skewed to the left. Mouse width was measured as the euclidean distance between the side laterals, and mouse length was measured as the euclidean distance between the nose and the tail base. Locomotion was defined as frames when the body center had a speed > 0.5 cm/s; motionless was defined as frames when the ears, body center, laterals, and tail base all had a speed ≤ 0.5 cm/s. To classify rearing, we constructed a random forest classifier in SimBA. 19,058 rearing frames from 35 video recordings of open-field behavior were extracted and manually annotated as rearing by three independent annotators. Supported and unsupported rearing behaviors were not differentiated. The start frame was defined as the frame in which the mouse lifted its forelimbs off the floor and extended its head upwards; the end frame was defined as the frame before the forelimbs made contact with the floor. The model was built with the following settings: n_estimators = 2,500, RF_criterion = entropy, RF_max_features = sqrt, RF_min_sample leaf = 2, and no oversampling or undersampling. 20% of the video frames were used for testing and the other 80% were used for training. The resulting classifier has a F1-score = 0.71, precision = 0.68, and recall = 0.74. The performance of this classifier was on par with those reported recently (Nilsson et al., 2020). The discrimination threshold was set at Pr = 0.31, and each instance of rearing had a minimum duration of 300 ms. Lastly, fine movement was defined as frames that did not fall into any of the categories mentioned above (i.e., locomotion, motionless, or rearing). Finally, example videos and the trained model are available on Github (https://github.com/saviochan/SimBA-OpenFieldArena) and Zenodo (https://zenodo.org/record/3964701#.XyB8yJ5KhPZ). The data generated by the analysis pipeline were processed using custom Python scripts. Codes are available online (https://github.com/saviochan/Python-Scripts/tree/master/OpenFieldArena_Behavior). Twenty-five different movement metrics were tracked. Event frequency, duration, and percent time spent were logged. ‘Light-period’ corresponds to 10 s of light delivery. ‘Pre-period’ and ‘post-period’ correspond to the 10 s epoch before and after light delivery, respectively. Fold changes were calculated by dividing the movement metric during light-period by that in pre-period.

To assess the relationship between the measured movement metrics, a correlation matrix was constructed from binned, time-series data. Rearing-motionless switch frequency and motionless-rearing switch frequency were excluded because of the low occurrence of events. Hierarchical clustering of movement metrics was performed in ClustVis (https://biit.cs.ut.ee/clustvis/) (Metsalu and Vilo, 2015). Twenty-five movement metrics were included in the analysis. Mice with targeted optogenetic stimulation of Foxp2^+^ neurons, Kcng4^+^ neurons, Npas1^+^ neurons, and PV^+^ neurons, were included. Both rows and columns were clustered using correlation distance and average linkage. Movement metrics were centered and scaled. A *K*-nearest neighbors classification algorithm was implemented in JASP to construct a decision boundary matrix. All variables were scaled to have a mean of 0 and a standard deviation of 1. The data were split as follows: 60% training, 20% testing, and 20% validation.

### Visualized *ex vivo* electrophysiology

Mice in the age range postnatal day 55–100 were anesthetized with a ketamine-xylazine mixture and perfused transcardially with ice-cold aCSF containing the following (in mM): 125 NaCl, 2.5 KCl, 1.25 NaH_2_PO_4_, 2.0 CaCl_2_, 1.0 MgCl_2_, 25 NaHCO_3_, and 12.5 glucose, bubbled continuously with carbogen (95% O_2_ and 5% CO_2_). The brains were rapidly removed, glued to the stage of a vibrating microtome (Leica Instrument), and immersed in ice-cold aCSF. Parasagittal slices containing the GPe were cut at a thickness of 240 μm and transferred to a holding chamber where they were submerged in aCSF at 37 °C for 30 min and returned to room temperature for recording. Slices were then transferred to a small-volume (∼0.5 ml) Delrin recording chamber that was mounted on a fixed-stage, upright microscope (Olympus). Neurons were visualized using differential interference contrast optics (Olympus), illuminated at 735 nm (Thorlabs), and imaged with a 60× water-immersion objective (Olympus) and a CCD camera (QImaging). Genetically-defined neurons were identified by somatic eGFP or tdTomato fluorescence examined under epifluorescence microscopy with a daylight (6,500 K) LED (Thorlabs) and appropriate filters (Semrock).

Recordings were made at room temperature (20–22 °C) with patch electrodes fabricated from capillary glass (Sutter Instrument) pulled on a Flaming-Brown puller (Sutter Instrument) and fire-polished with a microforge (Narishige) immediately before use. Pipette resistance was typically ∼3–5 MΩ. For cell-attached and current-clamp recordings, the internal solution consisted of the following (in mM): 135 KMeSO_4_, 10 Na_2_phosphocreatine, 5 KCl, 5 EGTA, 5 HEPES, 2 Mg_2_ATP, 0.5 CaCl_2_, and 0.5 Na_3_GTP, with pH adjusted to 7.25–7.30 with KOH. For voltage-clamp recordings of inhibitory postsynaptic currents (IPSCs), a high-chloride internal solution consisted of the following (in mM): 120 CsCl, 10 Na_2_phosphocreatine, 5 HEPES, 5 tetraethylammonium-Cl, 2 Mg_2_ATP, 1 QX314-Cl, 0.5 Na_3_GTP, 0.5 CaCl_2_, 0.25 EGTA, and 0.2% (wt/vol) biocytin, pH adjusted to 7.25–7.30 with CsOH and a low-chloride internal solution consisted of the following (in mM): 125 CsMeSO_3_, 10 Na_2_-phosphocreatine, 5 tetraethylammonium chloride, 5 QX-314 Cl, 5 HEPES-K, 5 EGTA-K, 2 Mg_2_ATP, 0.5 CaCl_2_, 0.5 Na_3_GTP, 0.2% (w/v) biocytin, with pH adjusted to 7.25–7.30 with CsOH. Stimulus generation and data acquisition were performed using an amplifier (Molecular Devices), a digitizer (Molecular Devices), and pClamp (Molecular Devices). For current-clamp recordings, the amplifier bridge circuit was adjusted to compensate for electrode resistance and was subsequently monitored. The signals were filtered at 1 kHz and digitized at 10 kHz. KMeSO_4_ and Na_2_-GTP were from ICN Biomedicals and Roche, respectively. All other reagents were obtained from Sigma-Aldrich.

For optogenetic experiments, blue excitation wavelength (peak, ∼450 nm) from two daylight (6,500 K) LEDs (Thorlabs) was delivered to the tissue slice from both a 60× water immersion objective and a 0.9 numerical aperture air condenser with the aid of 520 nm dichroic beamsplitters (Semrock). Light delivery was made at the site of electrophysiological recordings with a field of illumination of 500–700 µm in diameter. Paired-pulse optogenetic activation of terminals was at 20 Hz with a light duration of 2 ms. To extract the properties of spontaneous IPSCs in identified GPe neurons, an analysis similar to that described in (Higgs and Wilson, 2016) was used. Briefly, an autocorrelation of spontaneous IPSC times was computed, and for each periodic component detected, the periodic peaks were fitted with a model curve to estimate the presynaptic firing frequency, the coefficient of variation (CV) of presynaptic interspike intervals, and the synaptic success probability. The unitary IPSC amplitudes and waveforms were estimated by subdividing spontaneous IPSCs based on their timing with respect to the surrounding periodic input.

The electrophysiological data used in this study were expanded from a previously published dataset (Abecassis et al., 2020). Hierarchical clustering of electrophysiological attributes was performed in ClustVis (https://biit.cs.ut.ee/clustvis/) (Metsalu and Vilo, 2015). All cell-attached and whole-cell measurements were included in this analysis. Both rows and columns were clustered using correlation distance and average linkage. Electrophysiological measurements were centered and scaled. Principal component analysis of electrophysiological attributes was performed using the R programming language. t-SNE (*k* = 2) was performed in JASP using the Hartigan-Wong algorithm with the exact same dataset that was used for hierarchical clustering. All variables were scaled. Partial correlation networks based on Pearson coefficients of electrophysiological parameters were analyzed in JASP and Cytoscape (Shannon et al., 2003) with the CentiScaPe plug-in (Scardoni et al., 2009). The complete dataset for electrophysiological measurements of the intrinsic properties and an interactive web tool can be found online: https://icbi-georgetown.shinyapps.io/Neuro_interactive_analysis/.

### Drugs

*R*-CPP and NBQX disodium salt were obtained from Tocris. CGP55845 and QX314-Cl were obtained from Abcam. Na_3_GTP and tetrodotoxin were from Roche and Alomone Laboratories, respectively. Other reagents not listed above were from Sigma-Aldrich. Drugs were dissolved as stock solutions in either water or DMSO and aliquoted and frozen at –30 °C prior to use. Each drug was diluted to the appropriate concentrations by adding to the perfusate immediately before the experiment. The final concentration of DMSO in the perfusate was < 0.1%.

### Experimental design and statistical analyses

General graphing and statistical analyses were performed with MATLAB (MathWorks), Prism (GraphPad), JASP (https://jasp-stats.org) and the R environment (https://www.r-project.org). Custom analysis codes are available on GitHub (https://github.com/chanlab). Sample size (n value) is defined by the number of observations (i.e., neurons, sections, or mice). When percentages are presented, *n* values represent only positive observations. No statistical method was used to predetermine sample size. Data in the main text are presented as median values ± median absolute deviations (MADs) (Leys et al., 2013) as measures of central tendency and statistical dispersion, respectively. Box plots are used for graphic representation of population data unless stated otherwise (Krzywinski and Altman, 2014; Streit and Gehlenborg, 2014; Nuzzo, 2016). The central line represents the median, the box edges represent the interquartile ranges, and the whiskers represent 10–90^th^ percentiles. Normal distributions of data were not assumed. Individual data points were visualized for small sizes or to emphasize variability in the datasets. Non-parametric statistics were used throughout. Comparisons of unrelated samples were performed using a Mann–Whitney U test and comparisons of paired samples were performed using the Wilcoxon signed rank test. The Kruskal-Wallis test was used to compare multiple groups, followed by the *post hoc* Dunn test to assess pairwise differences while adjusting for the number of pairwise comparisons. The Spearman exact test was used for evaluating correlation between variables with a threshold (α) of 0.05 for significance. Unless < 0.0001 or > 0.99, exact P values (two-tailed) are reported in the text. To avoid arbitrary cutoffs and visual clutter, levels of significance are not included in the figures. Statistical significance is assessed at a significance level of 0.05, after applying the Bonferroni correction to adjust for the number of comparisons.

## Results

Recent single-cell transcriptomic analyses have revealed four distinct neuron clusters within the GPe (Saunders et al., 2018). In addition to the well-established Npas1^+^-Foxp2^+^ neurons (blue) that form a subclass of Npas1^+^ neurons, Npas1^+^-Nkx2.1^+^ neurons (cyan) are the other subclass of Npas1^+^ neurons (**Figure 1a**). Furthermore, PV^+^ neurons can be subdivided into two clusters (red and teal) with differential expression of Npr3, Kcng4, and Npy2r. Raw data showing the expression patterns of Npr3, Kcng4, and Npy2r can be found online (http://dropviz.org/?_state_id_=6c13807c8420bc74). The expression of these markers in the adult GPe is in agreement with results from *in situ* hybridization (**Figure 1b**). While sparse signals were found with Npr3 and Kcng4, the signals associated with Npy2r were widespread through the GPe. Here, we sought to examine whether Npr3-Cre (Daigle et al., 2018), Kcng4-Cre (Duan et al., 2014), and Npy2r-Cre (Chang et al., 2015) mice could provide genetic access to unique GPe subpopulations.

**Figure 1.**
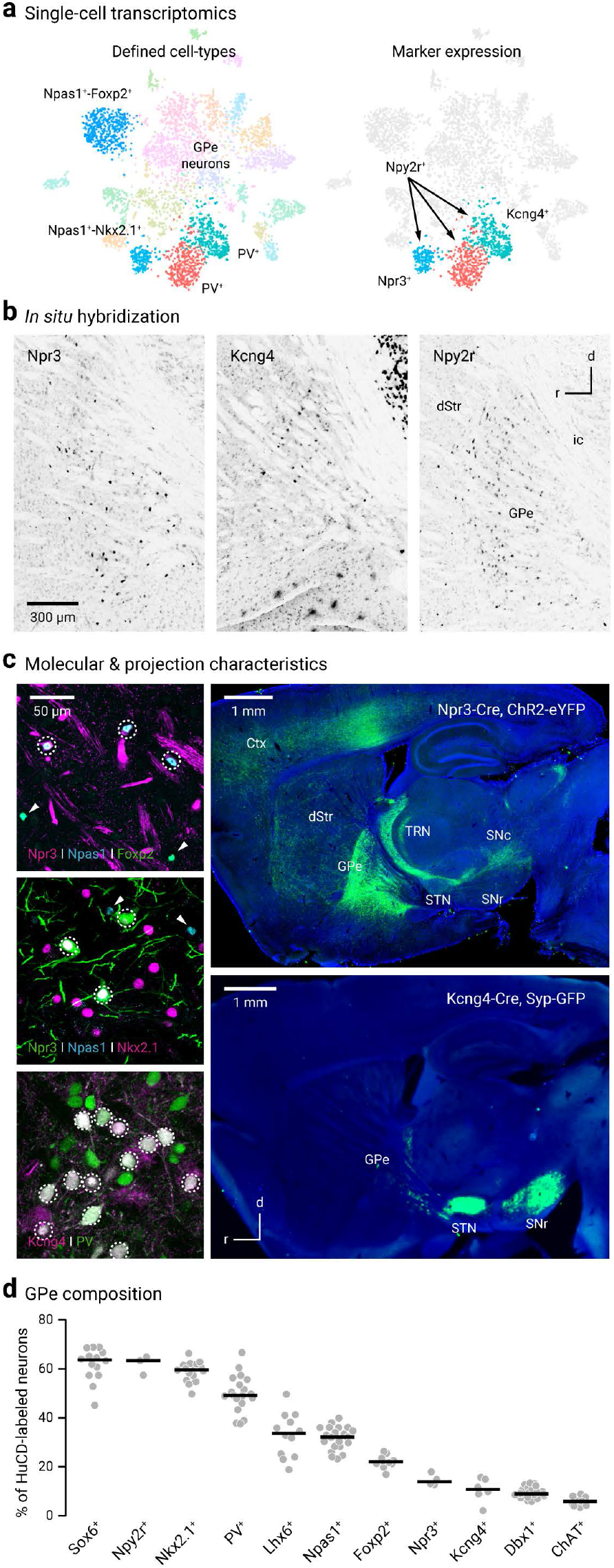
Novel markers and driver lines for interrogating GPe neurons. **a**. Left, Single-cell transcriptomic data identified four distinct clusters of GPe neurons; these include Npas1^+^-Foxp2^+^ (blue, 10 o’clock), Npas1^+^-Nkx2.1^+^(cyan, 7 o’clock), and two subpopulations of PV^+^ neurons (red and teal, 6 o’clock). Right, Npy2r is expressed in all non-Npas1^+^-Foxp2^+^ neurons; Npr3 and Kcng4 are expressed in a select subset of Npas1^+^ neurons and PV^+^ neurons, respectively. Figure adapted from (Saunders et al., 2018). Original expression data can be viewed at http://dropviz.org/?_state_id_=6c13807c8420bc74. **b**. Sagittal brain sections showing the *in situ* hybridization signals for Npr3 (left), Kcng4 (middle), Npy2r (right) in the GPe and neighboring areas. Data were adapted from Allen Brain Atlas. Brightness and contrast were adjusted. Raw data can be viewed and downloaded from http://mouse.brain-map.org. **c**. Top left, Coexpression (dotted circle) of Npas1 (blue) but not Foxp2 (green) in Npr3^+^ neurons (magenta). Npas1^+^-Foxp2^+^ (and non-Npr3^+^) neurons were visible in the same field (arrowheads). Middle left, Coexpression (dotted circle) of Npas1 (blue) and Nkx2.1 (magenta) in Npr3^+^ neurons (green). Npas1^+^ neurons that were neither Nkx2.1^+^ nor Npr3^+^ were visible in the same field (arrowheads). Bottom left, Coexpression (dotted circle) of PV (green) in Kcng4^+^ neurons (magenta). Top right, Npr3^+^ neurons share axonal projection patterns similar to that of Npas1^+^-Nkx2.1^+^; strong axonal projections to the cortex (Ctx), thalamic reticular nucleus (TRN), and substantia nigra pars compacta (SNc) were evident. Projections were assessed using viral delivery of ChR2-eYFP. Bottom right, Virally delivered synaptophysin-GFP in Kcng4-Cre mice revealed near-exclusive axonal projection from the external globus pallidus (GPe) to the subthalamic nucleus (STN) and substantia nigra pars reticulata (SNr). Abbreviations: dStr = dorsal striatum. **d**. Using HuCD as a neuronal marker, population data for the relative abundance of Gpe neuron markers were determined. Each circle represents a section. Medians and sample sizes are: Sox6^+^ (63.7%, 14), Npy2r^+^ (63.4%, 3), Nkx2.1^+^ (59.6%, 16), PV^+^ (49.2%, 19), Lhx6^+^ (33.7%, 12), Npas1^+^ (32.2%, 21), Foxp2^+^ (22.1%, 10), Npr3^+^ (14.0%, 4), Kcng4^+^ (10.8%, 6), Dbx1^+^ (9.0%, 22), ChAT^+^ (5.9%, 9). Data are ranked based on the median values as denoted by the thick lines. Parts of the data have been previously published.

### Novel driver lines capture distinct neuron populations

To study Npr3^+^ neurons, tdTomato-expressing (tdTomato^+^) neurons in Npr3-L-tdTomato mice were used. Npr3^+^ neurons were 14% of GPe neurons (14 ± 1%, *n* = 3 sections, 270 neurons). ∼90% of Npr3^+^ neurons were Npas1^+^ (89 ± 4%, *n* = 4 sections, 474 neurons), 74% were Nkx2.1^+^ (74 ± 0%, *n* = 3 sections, 169 neurons), and 2% were Foxp2^+^ (2 out of 170 neurons) (**Figure 1d**). To visualize the axonal projection pattern of Npr3^+^ neurons, ChR2-eYFP was transduced. This resulted in clear axonal arborization within the GPe, in addition to strong eYFP signals in the thalamic reticular nucleus, cortex, and substantia nigra pars compacta (**Figure 1c**). Our data are in line with the recent report that Lhx6^+^ GPe neurons strongly target compacta neurons (Evans et al., 2020). In summary, Npr3^+^ neurons had molecular and projectional properties that were completely consistent with those of Npas1^+^-Nkx2.1^+^ neurons (Abecassis et al., 2020). These findings further the idea that Npas1^+^-Nkx2.1^+^ neurons, along with the previously identified Npas1^+^-Foxp2^+^ neurons, are unique GPe neuron types and corroborate the findings in our recent study (Abecassis et al., 2020).

In addition to the robust expression in the thalamic reticular nucleus, *in situ* hybridization suggested that Kcng4 is expressed in a subset of neurons in the GPe. To further examine the molecular properties of Kcng4^+^ neurons, we performed immunohistological studies utilizing the Kcng4-L-tdTomato genetic cross. As shown in **Figure 1c**, the majority of Kcng4^+^ neurons (tdTomato^+^) express PV (90 ± 10%, *n* = 3 sections, 121 neurons) and they represent ∼30% (28 ± 8%, *n* = 3 sections, 121 neurons) of the PV^+^ population and 10% (10 ± 6%, *n* = 3 sections, 131 neurons) of the entirety of the GPe. These data are consistent with the single-cell transcriptomic data (Saunders et al., 2018) and show that Kcng4^+^ neurons are a subtype of PV^+^ neurons in the GPe. Using a Cre-inducible synaptophysin-GFP virus, we uncovered that Kcng4^+^ neurons send axonal projections to subthalamic nucleus (STN), internal globus pallidus, and substantia nigra pars reticulata. No additional projection targets were observed (**Figure 1c**).

To label Npy2r^+^ neurons, a similar genetic strategy was used (see Methods). tdTomato^+^ neurons from Npy2r-L-tdTomato mice were studied. As expected from the single-cell transcriptomic and *in situ* hybridization data, a large fraction (∼60%) of GPe neurons were tdTomato^+^. In addition to a population of GPe neurons, striosomes throughout the striatum were labeled in the Npy2r-L-tdTomato cross and appeared to be largely devoid of cell body labeling. The tdTomato^+^ neurons in the striatum were likely interneurons, based on their density (**Figure 2a**).

**Figure 2.**
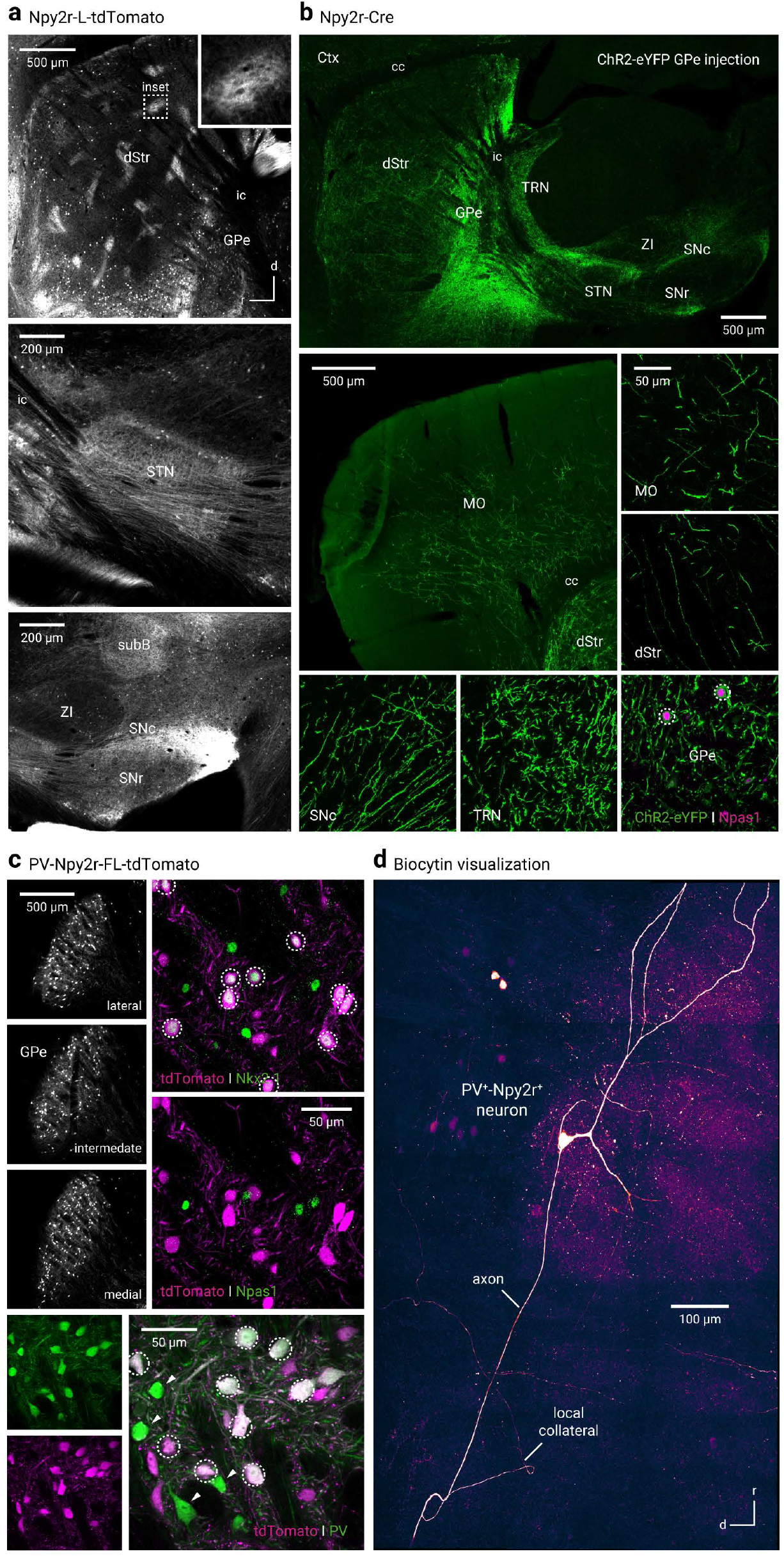
Npy2r-Cre mice capture a mixed population of Gpe neurons. **a**. Npy2r-L-tdTomato mice labeled neurons in the GPe and striosomes in the dorsal striatum (dStr) (top). Inset, a magnified view of a striosome. Prominent axons were apparent in the subthalamic nucleus (STN) (middle) and substantial nigra pars compacta (SNc) and reticulata (SNr) (bottom). **b**. Axonal projection patterns from Npy2r^+^ neurons are similar to that of Npas1^+^-Nkx2.1^+^ neurons and Npr3^+^ neurons. Only a low density of axons was observed in the subthalamic nucleus (STN) and substantial nigra pars reticulata (SNr). Lower right, ChR2-eYFP^+^ neurons were largely Npas1^+^. **c**. Top left, tdTomato-expressing (PV^+^-Npy2r^+^) neurons were observed throughout the GPe in PV-Npy2r-FL-tdTomato mice. PV^+^-Npy2r^+^ neurons were immunoreactive for Nkx2.1 (top right) but not for Npas1 (middle right). Bottom, most of PV^+^-Npy2r^+^ (tdTomato^+^) neurons were PV^+^. PV^+^-Npy2r^−^ (green) neurons were visible in the same field (arrowheads). **d**. A composite confocal micrograph showing a typical biocytin-filled PV^+^-Npy2r^+^ neuron. In addition to the main axon, local collaterals were observed. Abbreviations: cc = corpus callosum; dStr = dorsal striatum; ic = internal capsule; MO = motor cortex; SubB = subbrachial nucleus; TRN = thalamic reticular nucleus; ZI = zona incerta.

Though single-cell transcriptomic data suggest a broad expression profile of Npy2r among GPe neurons, whether the Npy2r-Cre mouse line would provide genetic access to selective neuron subtypes has not been tested. To this end, we transduced ChR2-eYFP to Cre^+^ neurons in Npy2r-Cre mice. This yielded axonal labeling patterns that were the most consistent with Npr3^+^ (aka Npas1^+^-Nkx2.1^+^) neurons (c.f. **Figure 1c**). In contrast, only sparse or weak axonal labeling was observed in canonical targets of PV^+^ neurons, i.e., STN and SNr (**Figure 2b**). The unexpected results prompted us to further examine the unusual properties of the Npy2r-Cre line. As roughly half of the tdTomato^+^ neurons (53 ± 8%, *n* = 11 sections, 1319 neurons) from Npy2r-L-tdTomato mice were PV^+^ (not shown), we decided to selectively study PV^+^-Npy2r^+^ neurons using the PV-Npy2r-FL-tdTomato genetic cross (**Figure 2c**). PV^+^-Npy2r^+^ neurons accounted for ∼65% (63 ± 4%, *n* = 11 sections, 1319 neurons) of the PV^+^ population and amounted to ∼35% (34 ± 1%, *n* = 3 sections, 392 neurons) of the entire GPe. The somatodendritic morphology of PV^+^-Npy2r^+^ neurons examined were also consistent with that of PV^+^ neurons (**Figure 2d**). To our surprise, a genetic PV-Npy2r-FL-tdTomato cross yielded tdTomato expression not only in PV^+^ neurons but also in a very small fraction (0.89%, *n* = 1 out of 112 neurons) of PV^−^ neurons (**Figure 2c**). One caveat of the FSF-LSL-tdTomato reporter strains is that they allow for very effective recombination, even for strains that have weak Cre activity. However, in strains that have high Cre activity, atypical expression patterns can be observed (Madisen et al., 2010).

### GPe neurons cluster into two electrophysiological groups

To study the electrophysiological properties of Npr3^+^ neurons, Kcng4^+^ neurons, Npy2r^+^ neurons, and PV^+^-Npy2r^+^ neurons, we performed *ex vivo* patch-clamp recordings using near-identical procedures described previously (Hernandez et al., 2015; Abecassis et al., 2020). To make sense of the results, we combined these data with an expanded version of a previously published dataset (Abecassis et al., 2020). Univariate scatter plots were used to emphasize the variability across samples. Corroborating the histological data, electrophysiological analysis showed that Npr3^+^ neurons and Kcng4^+^ neurons are a subset of the Npas1^+^ class and PV^+^ class, respectively. It is notable that Npr3^+^ neurons display properties that are indistinguishable from Npas1^+^-Lhx6^+^ neurons and Lhx6^+^_bright_ neurons. Apart from a strong tendency of Foxp2^+^ neurons and PV^+^ neurons to be on opposite ends of the spectrum, data are largely graded (**Figure 3**). Given that Kcng4 is an ion channel gene, it is surprising that the expression of Kcng4 did not confer unique electrophysiological properties.

**Figure 3.**
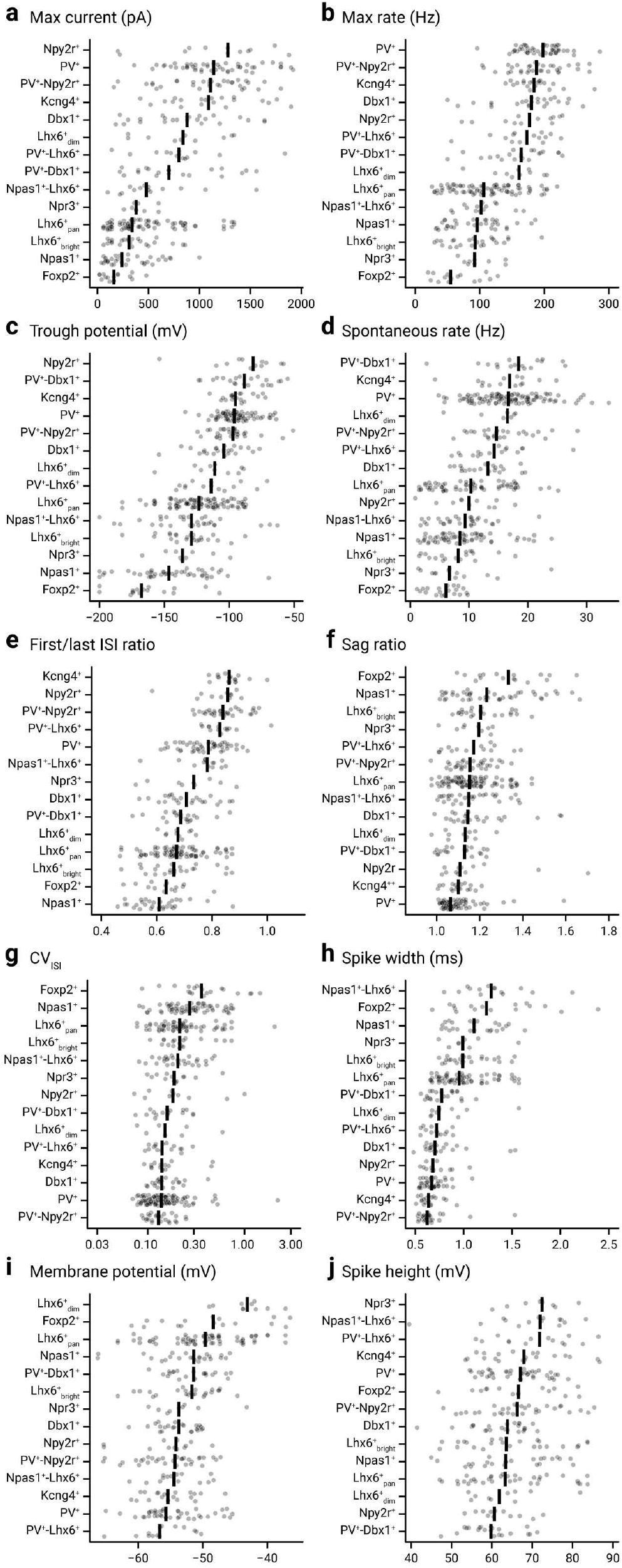
GPe neurons display graded electrophysiology attributes. Scatter plot summary of the electrophysiological properties of identified GPe neuron subtypes. Max current (**a**), max rate (**b**), trough potential (**c**), spontaneous rate (**d**), 1st/last ISI ratio (**e**), sag ratio (**f**), CV_ISI_ (**g**), spike width (**h**), membrane potential (**i**), and 1st spike height (**j**) are shown. See **Table 2** & **3** for a listing of median values, sample sizes, and statistical analysis of new neuron types, as compared to our recent publication (Abecassis et al., 2020), along with PV^+^ neurons and Npas1^+^ neurons for comparison. The data are ordered according to their median value as indicated by the thick vertical lines. Each cell is shown as a circle. A total of 329 neurons were included (n = Dbx1^+^: 24, Foxp2^+^: 16, Kcng4^+^: 20, Lhx6^+^_bright_: 24, Lhx6^+^_dim_: 10, Lhx6^+^_pan_: 68, Npas1^+^: 24, Npas1^+^-Lhx6^+^: 14, Npr3^+^: 9, Npy2r^+^: 12, PV^+^: 41, PV^+^-Dbx1^+^: 21, PV^+^-Lhx6^+^: 14, PV^+^-Npy2r^+^: 32) in this analysis. The complete dataset for electrophysiological measurements of the intrinsic properties and an interactive web tool can be found online: https://icbi-georgetown.shinyapps.io/Neuro_interactive_analysis/.

**Table 3.** Statistical analysis for electrophysiological characteristics.

|  | Kruskal-Wallis<sup>a</sup> | Pairwise comparisons<sup>b</sup> |  |  |  |  |  |  |  |  |  |  |  |  |  |
| --- | --- | --- | --- | --- | --- | --- | --- | --- | --- | --- | --- | --- | --- | --- | --- |
|  |  | Kcng4<sup>+</sup><br>vs<br>Npas1<sup>+</sup> | Kcng4<sup>+</sup><br>vs<br>Npas1<sup>+</sup>-Lhx6<sup>+</sup> | Kcng4<sup>+</sup><br>vs<br>Npr3<sup>+</sup> | Kcng4<sup>+</sup><br>vs<br>Npr2r<sup>+</sup> | Kcng4<sup>+</sup><br>vs<br>PV<sup>+</sup> | Kcng4<sup>+</sup><br>vs<br>PV<sup>+</sup>-Lhx6<sup>+</sup> | Kcng4<sup>+</sup><br>vs<br>PV<sup>+</sup>-<br>Npy2r<sup>+</sup> | Npas1<sup>+</sup><br>vs<br>Npas1<sup>+</sup>-Lhx6<sup>+</sup> | Npas1<sup>+</sup><br>vs<br>Npr3<sup>+</sup> | Npas1<sup>+</sup><br>vs<br>Npr2r<sup>+</sup> | Npas1<sup>+</sup><br>vs<br>PV<sup>+</sup> | Npas1<sup>+</sup><br>vs<br>PV<sup>+</sup>-Lhx6<sup>+</sup> | Npas1<sup>+</sup><br>vs<br>PV<sup>+</sup>-<br>Npy2r<sup>+</sup> | Npas1<sup>+</sup>-Lhx6<sup>+</sup><br>vs<br>Npr3<sup>+</sup> |
| Spontaneous activity (Hz) | **< 0.0001** | **0.0019** | **0.0037** | 0.014 | 0.38 | > 0.99 | > 0.99 | > 0.99 | > 0.99 | > 0.99 | > 0.99 | **< 0.0001** | 0.057 | **0.00010** | > 0.99 |
| CV<sub>ISI</sub> | **< 0.0001** | 0.014 | 0.97 | > 0.99 | > 0.99 | > 0.99 | > 0.99 | > 0.99 | > 0.99 | > 0.99 | > 0.99 | **< 0.0001** | 0.011 | **< 0.0001** | > 0.99 |
| Max rate (Hz) | **< 0.0001** | **< 0.0001** | **0.0014** | **0.0040** | > 0.99 | > 0.99 | > 0.99 | > 0.99 | > 0.99 | > 0.99 | **0.0029** | **< 0.0001** | 0.0059 | **< 0.0001** | > 0.99 |
| Max current (pA) | **< 0.0001** | **< 0.0001** | 0.046 | 0.019 | > 0.99 | > 0.99 | > 0.99 | > 0.99 | > 0.99 | > 0.99 | **< 0.0001** | **< 0.0001** | 0.022 | **< 0.0001** | > 0.99 |
| Membrane potential (mV) | 0.26 | > 0.99 | > 0.99 | > 0.99 | > 0.99 | > 0.99 | > 0.99 | > 0.99 | > 0.99 | > 0.99 | > 0.99 | > 0.99 | 0.38 | 0.79 | > 0.99 |
| Trough potential (mV) | **< 0.0001** | **< 0.0001** | **< 0.0001** | **0.0021** | > 0.99 | > 0.99 | 0.38 | > 0.99 | > 0.99 | > 0.99 | **< 0.0001** | **< 0.0001** | 0.072 | **< 0.0001** | > 0.99 |
| Sag ratio | **0.00011** | **0.00013** | > 0.99 | > 0.99 | > 0.99 | > 0.99 | > 0.99 | 0.99 | 0.022 | > 0.99 | 0.10 | **< 0.0001** | 0.19 | 0.099 | > 0.99 |
| First/last ISI ratio | **< 0.0001** | **< 0.0001** | 0.028 | **0.010** | > 0.99 | 0.019 | > 0.99 | > 0.99 | 0.40 | > 0.99 | **0.00036** | **0.0014** | **0.00031** | **< 0.0001** | > 0.99 |
| Spike height (mV) | 0.036 | > 0.99 | > 0.99 | > 0.99 | > 0.99 | > 0.99 | > 0.99 | > 0.99 | > 0.99 | > 0.99 | > 0.99 | > 0.99 | > 0.99 | > 0.99 | > 0.99 |
| Spike width (ms) | **< 0.0001** | **< 0.0001** | **< 0.0001** | **0.0016** | > 0.99 | > 0.99 | > 0.99 | > 0.99 | > 0.99 | > 0.99 | 0.0086 | **< 0.0001** | **0.0041** | **< 0.0001** | > 0.99 |

|  | Pairwise comparisons<sup>b</sup> |  |  |  |  |  |  |  |  |  |  |  |  |  |
| --- | --- | --- | --- | --- | --- | --- | --- | --- | --- | --- | --- | --- | --- | --- |
|  | Npas1<sup>+</sup>-Lhx6<sup>+</sup><br>vs<br>Npr2r<sup>+</sup> | Npas1<sup>+</sup>-Lhx6<sup>+</sup><br>vs<br>PV<sup>+</sup> | Npas1<sup>+</sup>-Lhx6<sup>+</sup><br>vs<br>PV<sup>+</sup>-Lhx6<sup>+</sup> | Npas1<sup>+</sup>-Lhx6<sup>+</sup><br>vs<br>PV<sup>+</sup>-Npy2r<sup>+</sup> | Npr3<sup>+</sup><br>vs<br>Npr2r<sup>+</sup> | Npr3<sup>+</sup><br>vs<br>PV<sup>+</sup> | Npr3<sup>+</sup><br>vs<br>PV<sup>+</sup>-Lhx6<sup>+</sup> | Npr3<sup>+</sup><br>vs<br>PV<sup>+</sup>-Npy2r<sup>+</sup> | Npy2r<sup>+</sup><br>vs<br>PV<sup>+</sup> | Npy2r<sup>+</sup><br>vs<br>PV<sup>+</sup>-Lhx6<sup>+</sup> | Npy2r<sup>+</sup><br>vs<br>PV<sup>+</sup>-<br>Npy2r<sup>+</sup> | PV<sup>+</sup><br>vs<br>PV<sup>+</sup>-Lhx6<sup>+</sup> | PV<sup>+</sup><br>vs<br>PV<sup>+</sup>-<br>Npy2r<sup>+</sup> | PV<sup>+</sup>-Lhx6<sup>+</sup><br>vs<br>PV<sup>+</sup>-Npy2r<sup>+</sup> |
| Spontaneous activity (Hz) | > 0.99 | **< 0.0001** | 0.081 | **0.00050** | > 0.99 | **< 0.0001** | 0.16 | **0.0050** | 0.029 | > 0.99 | 0.26 | > 0.99 | > 0.99 | > 0.99 |
| CV<sub>ISI</sub> | > 0.99 | **0.0043** | > 0.99 | 0.020 | > 0.99 | 0.98 | > 0.99 | 0.68 | > 0.99 | > 0.99 | > 0.99 | > 0.99 | > 0.99 | > 0.99 |
| Max rate (Hz) | 0.053 | **< 0.0001** | 0.11 | **< 0.0001** | 0.067 | **< 0.0001** | 0.13 | **0.00059** | > 0.99 | > 0.99 | > 0.99 | > 0.99 | > 0.99 | > 0.99 |
| Max current (pA) | 0.095 | **0.0018** | > 0.99 | 0.050 | 0.038 | **0.0011** | > 0.99 | 0.022 | > 0.99 | > 0.99 | > 0.99 | 0.30 | > 0.99 | > 0.99 |
| Membrane potential (mV) | > 0.99 | > 0.99 | > 0.99 | > 0.99 | > 0.99 | > 0.99 | > 0.99 | > 0.99 | > 0.99 | > 0.99 | > 0.99 | > 0.99 | > 0.99 | > 0.99 |
| Trough potential (mV) | **< 0.0001** | **< 0.0001** | > 0.99 | **0.00026** | **0.0015** | 0.0067 | > 0.99 | 0.016 | > 0.99 | 0.21 | > 0.99 | > 0.99 | > 0.99 | > 0.99 |
| Sag ratio | > 0.99 | > 0.99 | > 0.99 | > 0.99 | > 0.99 | > 0.99 | > 0.99 | > 0.99 | > 0.99 | > 0.99 | > 0.99 | > 0.99 | > 0.99 | > 0.99 |
| First/last ISI ratio | > 0.99 | > 0.99 | > 0.99 | 0.28 | 0.64 | > 0.99 | 0.79 | 0.090 | > 0.99 | > 0.99 | > 0.99 | > 0.99 | 0.28 | > 0.99 |
| Spike height (mV) | 0.47 | > 0.99 | > 0.99 | > 0.99 | 0.33 | 0.75 | > 0.99 | > 0.99 | > 0.99 | 0.30 | > 0.99 | 0.61 | > 0.99 | > 0.99 |
| Spike width (ms) | **0.0043** | **< 0.0001** | **0.0023** | **< 0.0001** | 0.27 | 0.0091 | 0.21 | **0.00038** | > 0.99 | > 0.99 | > 0.99 | > 0.99 | > 0.99 | > 0.99 |

Two-tailed P values are listed. <sup>a</sup>Kruskal–Wallis tests were performed to compare each characteristic across all groups. <sup>b</sup>Dunn tests were performed to compare each pair of neuron types. The bolded P values are significant when using a Bonferroni correction at a significance level of 0.05, adjusting for the fact that 10 characteristics were considered. Abbreviations: CV = coefficient of variation, ISI = interspike interval.

To more effectively visualize how Npr3^+^ neurons, Kcng4^+^ neurons, Npy2r^+^ neurons, and PV^+^-Npy2r^+^ neurons co-cluster with other known GPe neuron types, we performed principal component analysis (PCA) and a variety of clustering approaches on the previously published datasets (Abecassis et al., 2020) along with our newly collected dataset. Hierarchical clustering yielded two clusters that are largely consistent with the PV^+^ class and Npas1^+^ class. As expected, Npr3^+^ neurons co-clustered with the Npas1^+^ neuron class, while Kcng4^+^ neurons and PV^+^-Npy2r^+^ neurons co-clustered with the PV^+^ neuron class. As there was no *a priori* information on whether PV^+^ neurons can be subdivided based on their electrophysiological attributes, we clustered all identified PV^+^ neurons (*n* = 100 neurons) separate from the rest of the dataset (**Figure 4a**). t-SNE plots were used for visualizing high-dimensional data in a low-dimensional space. The relative distances between each marker represent similarities in the electrophysiological attributes. This analysis agreed with the PCA, in that two main clusters can be identified (**Figure 4b**). The PCA analysis shows that 45% of the total variance is explained by the first component and only 16% by the second component. The PCA biplot also shows the relative variance of each variable and their contribution to each principal component (**Figure 4c**). Maximal firing rate, maximal current, spike width, and trough potential made the most contributions to the first principal component. In contrast, the first action potential height and membrane potential contributed the least.

**Figure 4.**
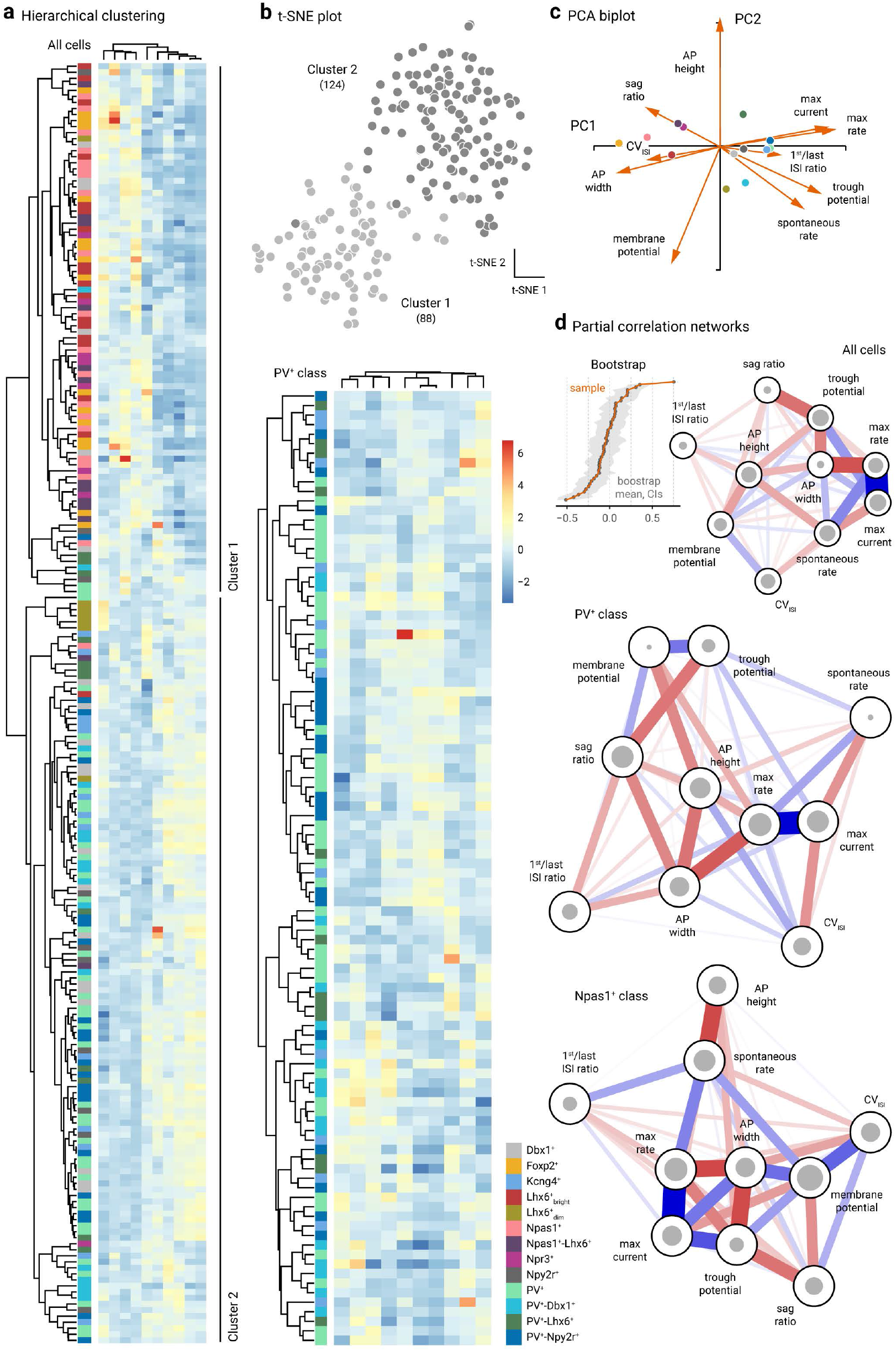
Clustering and network analysis of electrophysiological attributes. **a**. Left, Heatmap representation of electrical signatures of genetically-identified GPe neuron subtypes. Dendrograms show the order and distances of neuron clusters and their electrical characteristics. A total of 212 neurons (*n* = Dbx1^+^: 20, Foxp2^+^: 16, Kcng4^+^: 13, Lhx6^+^_bright_: 18, Lhx6^+^_dim_: 7, PV^+^: 38, PV^+^-Dbx1^+^: 16, PV^+^-Lhx6^+^: 12, PV^+^-Npy2r^+^: 21, Npas1^+^: 19, Npas1^+^-Lhx6^+^: 13, Npr3^+^: 8, Npy2r^+^: 10) were included in this analysis. Neurons with incomplete data were excluded from the analysis. Two main clusters were identified; cluster 1 contains 87 cells, cluster 2 contains 125 cells. Right, genetically-defined PV^+^ neurons are re-clustered. Scale bar applies to both panels. Purple-red-yellow colors were assigned for Npas1^+^ neurons, blue-green colors were assigned for PV^+^ neurons, and grays were used for mixed neuron types (i.e., Dbx1^+^ and Npy2r^+^). **b**. t-SNE analysis of the same data set as **a** yielded two distinct clusters. Cluster 1 (light gray) contains 88 cells, cluster 2 (dark gray) contains 124 cells. The membership assignment in cluster 1 and 2 for each neuron subtypes are as follows: Foxp2^+^ (100.0%, 0.0%), Npas1^+^ (95.2%, 4.8%), Lhx6^+^_bright_ (94.4%, 5.6%), Npr3^+^ (87.5%, 12.5%), Dbx1^+^ (30.0%, 70.0%), Npy2r^+^(30.0%, 70.0%), PV^+^-Lhx6^+^ (25.0%, 75.0%), Lhx6^+^_dim_ (14.3%, 85.7%), PV^+^-Dbx1^+^ (12.5%, 87.5%), PV^+^ (5.3%, 94.7%), PV^+^-Npy2r^+^ (4.8%, 95.2%), and Kcng4^+^ (0.0%, 100.0%). **c**. Principal component biplot showing the relative contributions of each electrophysiological attribute to PC1 and PC2. Markers represent the centroids for each cell type. The length of each vector is proportional to its variance with the cosine of the angle made by the vector with each axis indicating its contribution to that principal component. Highly correlated variables have similar directions; uncorrelated variables are perpendicular to each other. **d**. Partial correlation networks with a spring layout from all cells (top), PV^+^ neurons (middle), and Npas1^+^ neurons (bottom). Inset, bootstrap of edge stability. The size of the gray circles represents the connectivity of nodes to each other as measured by closeness centrality index. Edge thickness indicates the strength of correlation between nodes. Colors indicate the polarity of correlations. Blue indicates positive correlations; red indicates negative correlations.

To better understand the relationship between individual attributes and how they differed between the two neuron classes, we performed network analysis with the dataset. Subtle differences between the PV^+^ neuron and Npas1^+^ neuron networks were found (**Figure 4d**). A correlation between maximal firing rate, maximal current, and action potential width is consistent between the two networks and is conceivable as a function of congruent expression of voltage-gated sodium and potassium channels in these neurons (Hernandez et al., 2015).

### Markerless tracking effectively extracts behavioral dynamics

To investigate the roles of specific GPe populations in motor control, we analyzed the behavior of freely moving mice in an open field. We accomplished this by establishing a DeepLabCut (Mathis et al., 2018; Nath et al., 2019) model to track body kinematics (**Figure 5a**). The model was trained until the loss reached a plateau. This trained network had a test error of 1.13 pixels and a training error of 4.82 pixels. This error was negligible relative to the mouse size—the median mouse width and length were 65.2 ± 4.5 pixels and 152.8 ± 20.0 pixels, respectively. Speeds computed from the tracking data generated from DeepLabCut and Ethovision were tightly correlated, confirming the validity of our model (**Figure 5c**).

**Figure 5.**
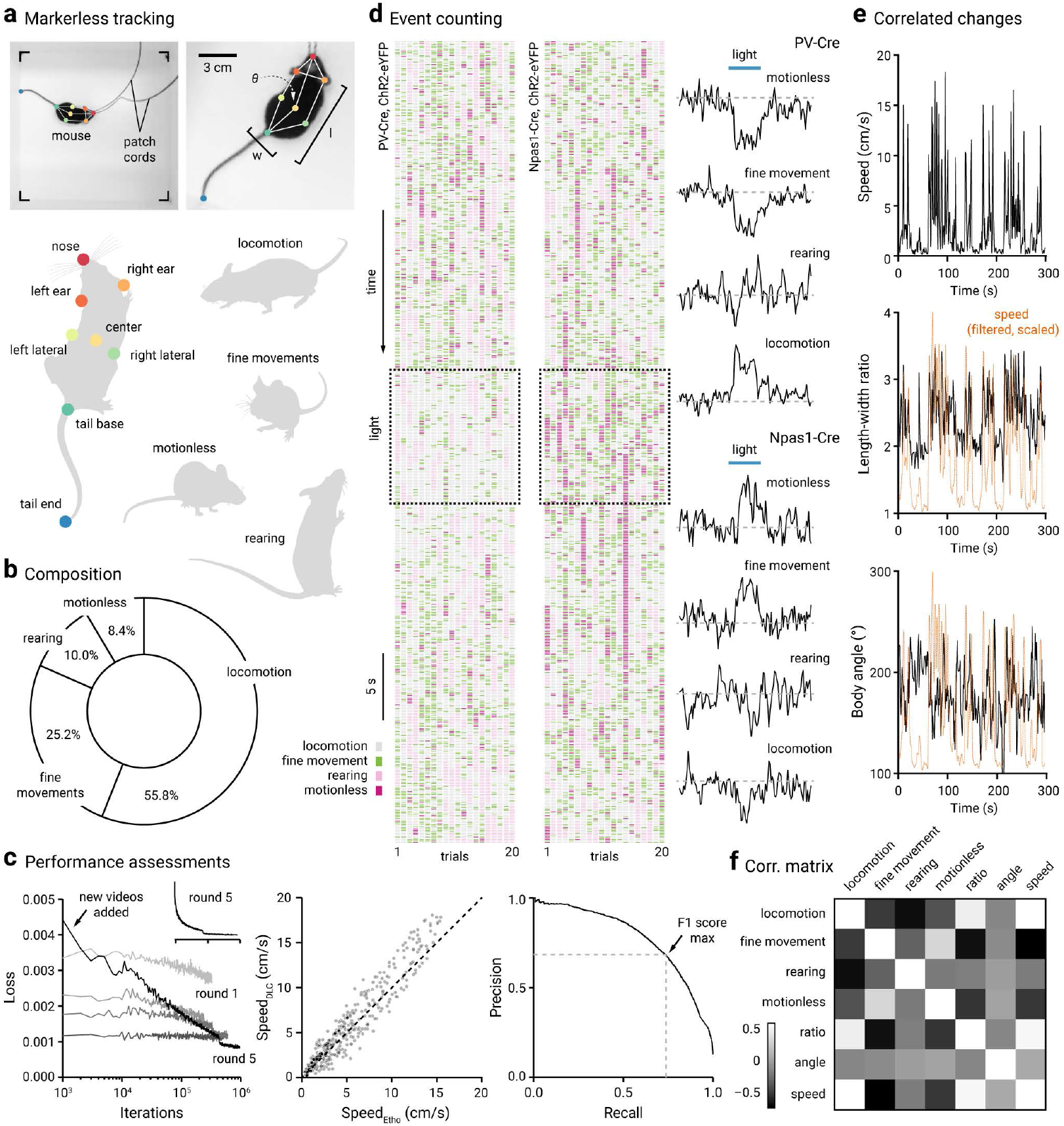
Markerless tracking and behavioral dynamics. **a**. Top left, A still frame showing a mouse exploring in an open-field arena. Patch cords were attached to the fiber implants for optogenetic interrogation. Color circles represent pose estimation markers. Top right, A magnified view of the same mouse shown on the left. Eight body parts were labeled and used for tracking body kinematics and motor patterns. Angle (*θ*), width (w), and length (l) were measured. Bottom, A schematized labeling configuration is shown. Right, Four discrete motor behavior motifs were extracted, namely, locomotion, fine movement, motionless, and rearing. **b**. The behavioral composition is shown as percent time spent. PV-Cre (n = 12) and Npas1-Cre (n = 16) mice virally transduced with ChR2-eYFP were included in this analysis. Mice were tethered to the patch cords but no light was delivered. **c**. Left, Learning curves of the DeepLabCut network over five rounds of training. Additional annotated frames were incorporated into the dataset in each round. An initial steep and monotonic decrease in loss was followed by a steady plateau. A large amount of new data was added to round 5, resulting in a high loss initially. The same data from round 5 were replotted on a linear scale as an inset (top right). Middle, An X-Y scatter plot comparing the speeds measured using DeepLabCut vs. Ethovision. Each marker is a frame. Diagonal line represents unity (i.e., x = y). Right, Precision-recall curve of the SimBA rearing classifier. Arrow indicates the maxima for the F1 score (0.71). **d**. Left & middle, Ethograms showing the motor dynamics of a PV-Cre mouse (left) and Npas1-Cre mouse (right) across 20 one-minute trials (columns). The beginning of each trial starts at the top of the plots. Square boxes show the time period in which light was delivered in the GPe. Rasters represent the occurrence of defined behavioral events: locomotion (gray), fine movement (green), rearing (light pink), and motionless (magenta). Each column is a single trial; twenty trials are shown for both PV-Cre and Npas1-Cre. Right, Population average changes in the event frequency of motionless, fine movement, rearing, and locomotion upon optogenetic stimulation (blue horizontal lines) of PV-Cre (n = 12, top) and Npas1-Cre (n = 16, bottom) mice transduced with ChR2-eYFP. **e**. Top to bottom, Correlated changes in speed, length-width ratio, and body angle of an example mouse across time. Speed data (orange dotted lines) were filtered and scaled to facilitate comparisons. **f**. A correlation matrix constructed using time series data from both pre-period and light-period illustrating moment-to-moment covariance of the different movement metrics. The event frequencies of the four behaviors, body angle, length-width ratio, and speed were used in this comparison. PV-Cre and Npas1-Cre mice (virally transduced with ChR2-eYFP) were included in this analysis (n = 28). Light colors indicate positive correlations; dark colors indicate negative correlations.

To understand the motor patterns of mice in an open field, we defined ‘locomotion’, ‘fine movement’, and ‘motionless’ based on the movements of body parts (**Figure 5a**). To describe the full repertoire of motor behavior, we captured ‘rearing’ in SimBA (Nilsson et al., 2020), which used markerless tracking data in conjunction with manual annotation to generate predictive classifiers. This classifier has a F1-score = 0.71, precision = 0.68, and recall = 0.74 (**Figure 5c**). Using this machine learning pipeline, we were able to effectively quantify behavioral events of mice in the open field in much greater detail and detect more complex, yet subtle, changes in body kinematics than before (**supplemental video**). We quantified the four different behaviors in PV-Cre and Npas1-Cre mice (*n* = 28 mice). At the population level, time spent in each category was: 55.8% in locomotion, 25.2% in fine movement, 10.0% in rearing, and 8.4% in motionless (**Figure 5b**).

Our previous studies established that PV^+^ neurons and Npas1^+^ neurons play causal but opposite roles in regulating locomotion—PV^+^ neurons are movement-promoting and Npas1^+^ neurons are movement-suppressing (Glajch et al., 2016; Pamukcu et al., 2020). Consistent with our earlier observations, our analysis here showed that mice exhibited more mobile behavior (i.e., locomotion) when PV^+^ neurons were optogenetically stimulated and more immobile behavior (i.e., fine movement and motionless) when Npas1^+^ neurons were optogenetically stimulated. On the other hand, rearing was not consistently altered by optogenetic perturbations of either PV^+^ neurons or Npas1^+^ neurons (**Figure 5d**). This provides a proof-of-concept that the network can be used effectively to extract body kinematics in mice. To monitor the orientation and posture of the mice, we measured the body angle and length-width ratio, respectively (**Figure 5e**). A correlation matrix was constructed to visualize the relationships between different movement metrics (**Figure 5f**). Spearman rank correlation tests between pairs of variables showed that 10 of the 21 pairs were correlated after the Bonferroni correction: locomotion was correlated with fine movement, rearing, body ratio, and speed; fine movement was correlated with motionless, body ratio, and speed; motionless was correlated with body ratio and speed; and body ratio was correlated with speed. In particular, the average event counts of fine movement and motionless were positively correlated with each other. Locomotion was associated with a high length-width ratio and a high speed, indicating that the mouse assumed an elongated form during locomotion. Fine movement and motionless were associated with a low length-width ratio and a low speed, which is indicative of a hunched and retracted posture. Similarly, body angle correlated with moment-to-moment changes in speed and body ratio (**Figure 5e**).

### Optogenetic manipulations induce cell-specific motor patterns

As a large number of parameters (a total of 25) including movement sequences were tracked, we examined the relationships between the measured parameters by constructing a correlation matrix for both PV-Cre and Npas1-Cre mice that were used in this study (*n* = 28). We found strong correlations between a large number of variable-pairs (163 from 253), assessed via Spearman rank correlation tests after the Bonferroni correction. As mice have low incidences of transitioning between rearing and motionless, rearing-motionless and motionless-rearing switch frequencies were excluded from this analysis. At this point, it is unclear if that was hardwired in the motor program or if it was constrained by body mechanics. Data from pre- and light-periods were combined, as no marked differences in the relationships between movement metrics were noted (not shown). A condensed version of the correlation matrix that contains the most relevant metrics are presented in **Figure 6a**. For example, percent time spent on locomotion positively correlated (blue) with speed but negatively correlated (brown) with percent time spent on fine movement and motionless. To effectively visualize the behavioral responses in mice when PV^+^ neurons and Npas1^+^ neurons were optogenetically stimulated, we performed a PCA. As shown in **Figure 6a**, mice showed very similar behavior during the pre-period. In contrast, they showed divergent responses when PV^+^ neurons and Npas1^+^ neurons were optogenetically stimulated during the light-period.

**Figure 6.**
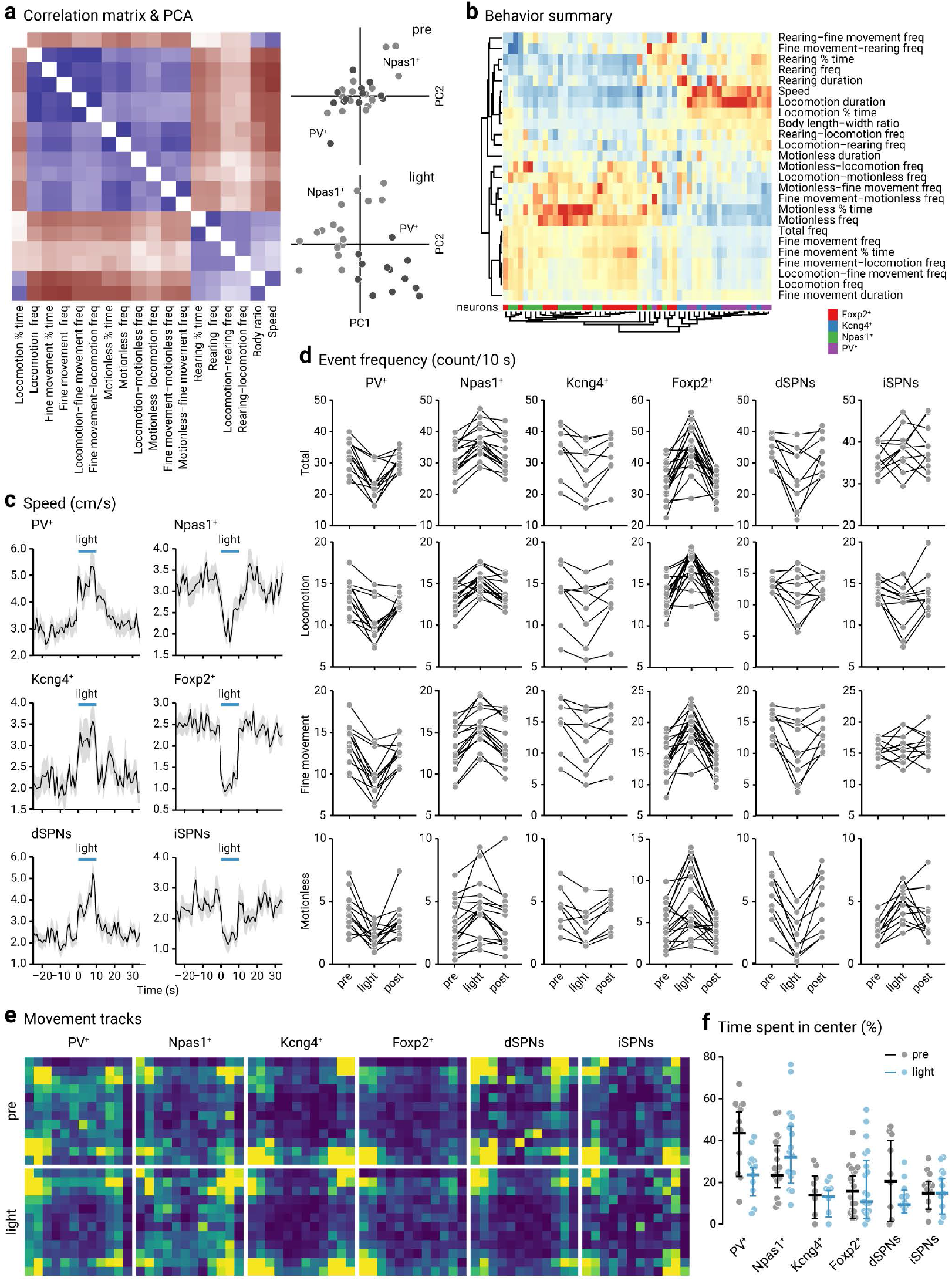
Motor patterns of mice with cell-specific optogenetic stimulation. **a**. Left, A correlation matrix constructed using data from PV-Cre (n = 12) and Npas1-Cre (n = 16) mice transduced with ChR2-eYFP. Eighteen parameters were included in this matrix. Blue colors indicate positive correlations, brown colors indicate negative correlations. Right, PCA plots showing the distributions of PV-Cre (dark gray, n = 12) and Npas1-Cre (light gray, n = 16) mice (transduced with ChR2-eYFP) in pre-period (top) and light-period (bottom). Each marker represents a mouse. Twenty three movement metrics were used in this analysis. **b**. A heatmap summarizing motor responses of mice to optogenetic stimulation of genetically-defined neurons in the GPe. Twenty-five movement metrics were measured to fully capture the behavioral structures. Each of the 25 rows represents the fold change of movement metrics. Warm colors (red) represent positive changes; cool colors (blue) represent negative changes. Rows and columns were sorted using hierarchical clustering. Dendrograms are divided into two main arms; metrics on the upper arm are negatively correlated with ‘total frequency’, while metrics on the lower arm are positively correlated with ‘total frequency’. Each column is a mouse; 54 mice were used in this analysis. The neurons of interest are PV^+^ (purple), Npas1^+^ (green), Kcng4^+^ (blue), and Foxp2^+^ (red) neurons. **c**. Plots showing the speed of mice with optogenetic stimulation in genetically-defined neurons in the GPe. Blue horizontal lines indicate the timing of light delivery. Thick lines indicate average, gray areas indicate standard error of the means. **d**. Slopegraphs showing event frequency (counts per 10 s) for total, locomotion, fine movement, and motionless in mice and the effect with optogenetic stimulation of selective neuron types. Each connected line represents a mouse: PV^+^ neurons (n = 12), Npas1^+^ neurons (n = 16), Kcng4^+^ neurons (n = 8), Foxp2^+^ neurons (n = 18), dSPNs (n = 9), and iSPNs (n = 11). **e**. Heatmaps showing the relative cumulative time that mice spent in each position following optogenetic stimulation of genetically-defined GPe neurons. Yellow represents more time spent; blue represents less time spent. Each heatmap was scaled to min and max and does not represent the net locomotor activity of mice. **f**.A univariate scatter plot summarizing the amount of time mice spent in the center of the area during pre- (gray) and light-period (blue). PV^+^ neurons (n = 12), Npas1^+^ neurons (n = 16), Kcng4^+^ neurons (n = 8), Foxp2^+^ neurons (n = 18), dSPNs (n = 9), and iSPNs (n = 11).

**Figure 6b** is a heatmap representation of changes in movement metrics upon optogenetic stimulation of genetically-defined GPe neurons. Mice transduced with ChR2-eYFP in Foxp2^+^ neurons, Kcng4^+^ neurons, Npas1^+^ neurons, and PV^+^ neurons were included in this analysis. Hierarchical clustering revealed coordinated changes in behavioral dynamics across these test subjects. In particular, the stimulation of Foxp2^+^ neurons and Kcng4^+^ neurons led to changes that were consistent with those produced by the stimulation of Npas1^+^ neurons and PV^+^ neurons, respectively. Unlike the ethograms in **Figure 5d** that are purely event-counting across time, the heatmap in **Figure 6b** shows the fold changes of all 25 movement metrics in response to optogenetic stimulation. The most striking effects were the changes in speed, locomotion duration, percent time in motionless, and motionless frequency. The raw data for speed is shown in **Figure 6c**. To contextualize the magnitude of the observed effect, direct- and indirect-pathway striatal projection neurons (dSPNs and iSPNs, respectively) were included for comparison. **Figure 6d** shows the changes in total, locomotion, fine movement, and motionless frequencies across individual mice. The subtle differences between mouse lines made it difficult to make arguments about quantitative differences. With this caveat in mind, it is nevertheless surprising to see that optogenetic stimulation of Kcng4^+^ neurons produced statistically indistinguishable effects on locomotor speed as PV^+^ neurons. This result was not expected as Kcng4^+^ neurons only account for ∼10% of the entirety of the GPe. Similarly, optogenetic stimulation of Foxp2^+^ neurons produced near-identical effects on locomotor speed as Npas1^+^ neurons (**Table 4**), while they only account for ∼25% of the entirety of the GPe (**Figure 1d**).

**Table 4.** Summary of optogenetic effects on behavior metrics.

|  | PV+ neurons, ChR2 (n = 12) |  | Npas1+ neurons, ChR2 (n = 16) |  | Kcng4+ neurons, ChR2 (n = 8) |  | Foxp2+ neurons, ChR2 (n = 18) |  | STN neurons, ACR2 (n = 10) |  | dSPNs, ChR2 (n = 9) |  | iSPNs, ChR2 (n = 11) |  |
| --- | --- | --- | --- | --- | --- | --- | --- | --- | --- | --- | --- | --- | --- | --- |
|  | Fold change | P value<sup>c</sup> | Fold change | P value | Fold change | P value | Fold change | P value | Fold change | P value | Fold change | P value | Fold change | P value |
| Speed (cm/s) | 1.53 ± 0.31 | 0.00098 | 0.81 ± 0.09 | **0.000031** | 1.52 ± 0.33 | 0.0078 | 0.36 ± 0.12 | **0.0000076** | 1.68 ± 0.15 | 0.0020 | 1.9 ± 0.25 | 0.0039 | 0.52 ± 0.09 | 0.00098 |
| Body length-width ratio | 1.1 ± 0.05 | 0.00049 | 1.0 ± 0.05 | **0.00015** | 1.16 ± 0.08 | 0.023 | 0.88 ± 0.06 | **0.000023** | 1.19 ± 0.03 | 0.0020 | 1.14 ± 0.11 | 0.0039 | 0.97 ± 0.04 | 0.083 |
| Total frequency<sup>a</sup> | 0.71 ± 0.12 | 0.00049 | 1.08 ± 0.08 | **0.000061** | 0.79 ± 0.06 | 0.0078 | 1.36 ± 0.17 | **0.000015** | 0.68 ± 0.06 | 0.0020 | 0.76 ± 0.16 | 0.0039 | 1.12 ± 0.09 | 0.31 |
| Locomotion frequency | 0.78 ± 0.12 | 0.00049 | 1.1 ± 0.04 | **0.000092** | 0.82 ± 0.1 | 0.023 | 1.32 ± 0.18 | **0.000076** | 0.71 ± 0.05 | 0.0020 | 0.91 ± 0.18 | 0.20 | 0.9 ± 0.04 | 0.0098 |
| Fine movement frequency | 0.69 ± 0.12 | 0.00049 | 1.06 ± 0.1 | **0.000061** | 0.77 ± 0.13 | 0.0078 | 1.35 ± 0.21 | **0.000015** | 0.63 ± 0.08 | 0.0020 | 0.82 ± 0.14 | 0.0039 | 1.04 ± 0.08 | 0.81 |
| Rearing frequency | 0.91 ± 0.2 | 0.15 | 0.87 ± 0.18 | 0.026 | 0.98 ± 0.13 | 0.56 | 0.68 ± 0.31 | 0.00092 | 1.56 ± 0.33 | 0.0039 | 0.97 ± 0.32 | 0.97 | 0.29 ± 0.11 | 0.00098 |
| Motionless frequency | 0.5 ± 0.18 | 0.0015 | 1.03 ± 0.46 | 0.0011 | 0.67 ± 0.14 | 0.016 | 1.77 ± 0.34 | 0.0019 | 0.51 ± 0.13 | 0.0059 | 0.33 ± 0.14 | 0.0039 | 1.88 ± 0.34 | 0.00098 |
| Locomotion % time | 1.24 ± 0.09 | 0.0024 | 0.99 ± 0.07 | 0.0013 | 1.11 ± 0.04 | 0.0078 | 0.86 ± 0.06 | 0.0023 | 1.18 ± 0.13 | 0.0059 | 1.37 ± 0.12 | 0.0039 | 0.66 ± 0.1 | 0.0029 |
| Fine movement % time | 0.66 ± 0.14 | 0.00049 | 1.09 ± 0.14 | **0.000031** | 0.74 ± 0.16 | 0.0078 | 1.51 ± 0.32 | **0.000038** | 0.53 ± 0.12 | 0.0020 | 0.79 ± 0.13 | 0.0039 | 1.03 ± 0.08 | 0.64 |
| Rearing % time | 1.02 ± 0.25 | 0.91 | 0.82 ± 0.16 | 0.013 | 1.21 ± 0.14 | 0.023 | 0.54 ± 0.25 | 0.0010 | 1.42 ± 0.25 | 0.020 | 1.11 ± 0.57 | 0.65 | 0.39 ± 0.18 | 0.00098 |
| Motionless % time | 0.41 ± 0.15 | 0.0024 | 1.08 ± 0.49 | 0.0063 | 0.63 ± 0.2 | 0.0078 | 1.43 ± 0.53 | 0.060 | 0.5 ± 0.19 | 0.020 | 0.18 ± 0.1 | 0.0039 | 2.84 ± 0.34 | 0.00098 |
| Locomotion duration<sup>b</sup> | 1.83 ± 0.36 | 0.00049 | 0.91 ± 0.05 | **0.000031** | 1.38 ± 0.17 | 0.0078 | 0.57 ± 0.08 | **0.000015** | 1.58 ± 0.21 | 0.0020 | 1.7 ± 0.41 | 0.0039 | 0.95 ± 0.23 | 0.21 |
| Fine movement duration | 0.96 ± 0.03 | 0.016 | 1.01 ± 0.06 | 0.011 | 0.94 ± 0.04 | 0.039 | 1.16 ± 0.1 | 0.0010 | 0.85 ± 0.08 | 0.0098 | 0.93 ± 0.07 | 0.027 | 0.94 ± 0.05 | 0.019 |
| Rearing duration | 1.17 ± 0.13 | 0.11 | 0.99 ± 0.11 | 0.074 | 1.34 ± 0.49 | 0.31 | 0.7 ± 0.12 | 0.0066 | 0.92 ± 0.12 | 0.16 | 0.84 ± 0.29 | 0.82 | 1.28 ± 0.56 | 0.43 |
| Motionless duration | 0.78 ± 0.1 | 0.0093 | 0.95 ± 0.09 | 0.86 | 0.95 ± 0.13 | 0.38 | 0.88 ± 0.11 | 0.067 | 1.0 ± 0.28 | 0.63 | 0.61 ± 0.08 | 0.0039 | 1.67 ± 0.32 | 0.0020 |
| Locomotion-fine movement frequency | 0.77 ± 0.14 | 0.00049 | 1.08 ± 0.08 | **0.000061** | 0.79 ± 0.17 | 0.039 | 1.32 ± 0.26 | 0.00033 | 0.62 ± 0.04 | 0.0020 | 1.01 ± 0.24 | 0.50 | 0.72 ± 0.13 | 0.0020 |
| Locomotion-rearing frequency | 1.02 ± 0.13 | 0.93 | 1.0 ± 0.18 | 0.75 | 0.87 ± 0.11 | 0.054 | 0.76 ± 0.11 | 0.083 | 1.43 ± 0.07 | 0.0020 | 1.14 ± 0.13 | 0.31 | 0.57 ± 0.29 | 0.031 |
| Locomotion-motionless frequency | 0.7 ± 0.21 | 0.034 | 1.06 ± 0.31 | 0.021 | 0.88 ± 0.25 | 0.64 | 1.37 ± 0.35 | 0.0019 | 0.73 ± 0.21 | 0.29 | 0.77 ± 0.06 | 0.0039 | 1.38 ± 0.35 | 0.0088 |
| Fine movement-locomotion frequency | 0.77 ± 0.11 | 0.00049 | 1.06 ± 0.11 | **0.00015** | 0.79 ± 0.16 | 0.039 | 1.35 ± 0.24 | **0.00011** | 0.64 ± 0.08 | 0.0020 | 1.02 ± 0.3 | 0.48 | 0.8 ± 0.12 | 0.0020 |
| Fine movement-rearing frequency | 0.95 ± 0.08 | 0.12 | 1.0 ± 0.22 | 0.39 | 0.99 ± 0.08 | > 0.99 | 0.96 ± 0.16 | 0.54 | 0.97 ± 0.2 | 0.47 | 0.86 ± 0.46 | 0.44 | 0.0 ± 0.0 | > 0.99 |
| Fine movement-motionless frequency | 0.71 ± 0.15 | 0.034 | 1.06 ± 0.2 | 0.034 | 0.72 ± 0.12 | 0.078 | 1.31 ± 0.36 | 0.048 | 0.6 ± 0.11 | 0.063 | 0.45 ± 0.11 | 0.0039 | 1.56 ± 0.36 | 0.00098 |
| Rearing-locomotion frequency | 1.09 ± 0.16 | 0.83 | 1.03 ± 0.18 | 0.9 | 0.95 ± 0.16 | 0.74 | 0.81 ± 0.19 | 0.086 | 1.17 ± 0.06 | 0.027 | 1.07 ± 0.18 | 0.81 | 0.67 ± 0.17 | 0.0078 |
| Rearing-fine movement frequency | 1.0 ± 0.0 | 0.25 | 1.0 ± 0.15 | 0.57 | 0.95 ± 0.06 | 0.13 | 0.9 ± 0.13 | 0.49 | 0.78 ± 0.28 | 0.13 | 1.0 ± 0.14 | 0.63 | 1.0 ± 0.33 | > 0.99 |
| Motionless-locomotion frequency | 0.71 ± 0.13 | 0.077 | 1.01 ± 0.33 | 0.0082 | 0.92 ± 0.17 | 0.74 | 1.31 ± 0.33 | 0.054 | 0.78 ± 0.13 | 0.027 | 0.71 ± 0.07 | 0.0078 | 1.42 ± 0.27 | 0.0059 |
| Motionless-fine movement frequency | 0.72 ± 0.12 | 0.027 | 0.96 ± 0.39 | 0.074 | 0.71 ± 0.09 | 0.023 | 1.36 ± 0.25 | 0.027 | 0.72 ± 0.31 | 0.29 | 0.43 ± 0.11 | 0.0039 | 1.67 ± 0.35 | 0.00098 |

<sup>a</sup>Unit for all frequencies is count/10 s. <sup>b</sup>Unit for all durations is seconds. <sup>c</sup>Wilcoxon test; two-tailed exact P values are shown. The bolded P values are significant when using a Bonferroni correction at a significance level of 0.05.

Given Kcng4^+^ neurons are ∼30% (28 ± 8%, *n* = 3 sections, 121 neurons) of PV^+^ neurons and Foxp2^+^ neurons are ∼60% (57 ± 4%, *n* = 6 sections, 566 neurons) of Npas1^+^ neurons, our results raised a question of whether PV^+^-Kcng4^−^ neurons and Npas1^+^-Foxp2^−^ neurons could be involved in non-motor function. Close examination of the movement tracks revealed that optogenetic stimulation of PV^+^ neurons and Npas1^+^ neurons led to changes in the anxiety-like behavior, as measured by their center-edge preference in the open-field arena. As shown in **Figure 6e** & **f**, PV^+^ neuron stimulation decreased the time spent at the center, while Npas1^+^ neuron stimulation increased it (PV^+^ pre: 47.3 ± 10.3%, PV^+^ light: 27.4 ± 5.4%, *n* = 12, P = 0.0034; Npas1^+^ pre: 27.3 ± 10.4%, Npas1^+^ light: 35.7 ± 12.3%, *n* = 16, P = 0.0060). These effects were highly cell type-specific as they were not observed with the stimulation of Kcng4^+^ neurons, Foxp2^+^ neurons, dSPNs, or iSPNs and thus were dissociable from the locomotor effects (Kcng4^+^ pre: 17.7 ± 7.9%, Kcng4^+^ light: 16.9 ± 4.1%, *n* = 8, P = 0.38; Foxp2^+^ pre: 11.5 ± 10.7%, Foxp2^+^ light: 10.4 ± 8.7%, *n* = 18, P = 0.30; dSPN pre: 24.1 ± 19.0%, dSPN light: 13.1 ± 4.1%, *n* = 9, P = 0.10; iSPN pre: 18.6 ± 7.2%, iSPN light: 18.6 ± 9.6%, *n* = 11, P = 0.90).

### Combinatorial movement metrics infers behavioral states

To visualize how each neuron subtype would be classified based on their motor effects upon optogenetic stimulation, we constructed a decision boundary matrix using a *k*-nearest neighbors classification algorithm (**Figure 7a**). As supported by the receiver operating characteristic (ROC) curves in **Figure 7b**, each neuron type studied here gave rise to unique behavior structures. This analysis showed that a number of neuron types occupied a unique parameter space as delineated by the decision boundaries visualized (e.g., **Figure 7c**). As shown in the relationship between locomotion frequency and the percent time in locomotion (**Figure 7c**), PV^+^ neurons and Npas1^+^ neurons were on opposite ends of the spectrum. This suggests that PV^+^ neurons and Npas1^+^ neurons work against each other. On the other hand, PV^+^ neurons, Kcng4^+^ neurons, dSPNs, and STN neurons (transduced with GtACR2, an inhibitory opsin) were close to one another, suggesting that these neurons work either together or converge on a common downstream target to mediate the same action. This idea is supported by experimental evidence that PV^+^ neurons and Kcng4^+^ neurons send inhibitory signals to the substantia nigra pars reticulata, which receive direct inhibitory inputs from dSPNs (Grofova, 1975; Araki et al., 1985; Bolam et al., 1993; Mink, 1996; Smith et al., 1998; Connelly et al., 2010; Cui et al., 2020). In contrast, iSPNs that are known to heavily target PV^+^ neurons (but not PV^−^ neurons) (Yuan et al., 2017) are on the lower-left corner of the plot. This analysis reinforced the notion that the differences in the tracked motor behaviors were in fact cell-specific and not simply esoteric fluctuations.

**Figure 7.**
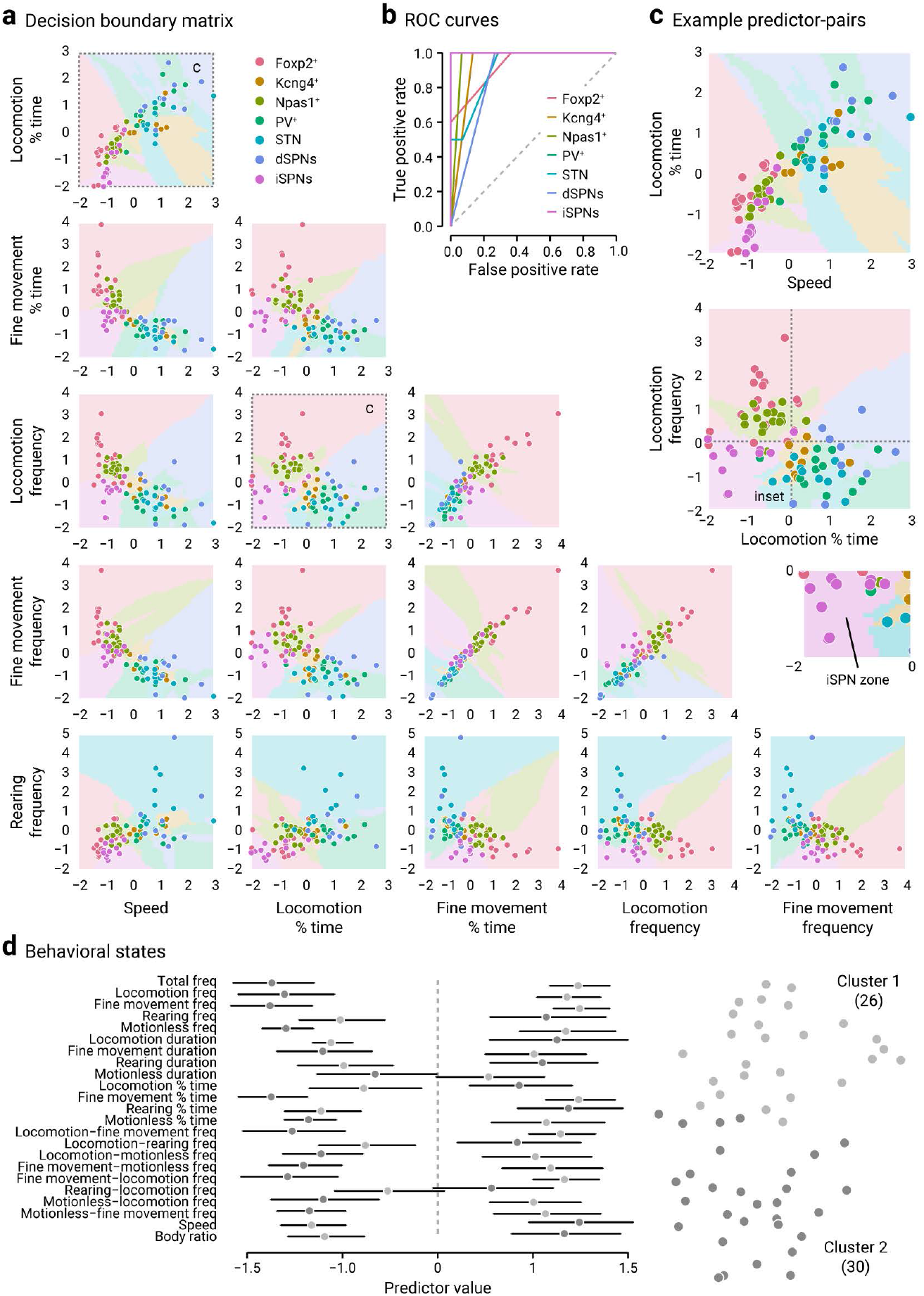
Optogenetic manipulations produced cell-specific motor patterns. **a**. A decision boundary matrix visualizing the relationships between movement metrics and how they would be classified if predicted through the current model. Boundaries between classes are visualized. All variables were scaled to have a mean of 0 and a standard deviation of 1. **b**. Receiver operating characteristic (ROC) curves for different cell types are shown. Classifiers that give curves closer to the top-left corner indicate a better performance; the closer the curve comes to the diagonal of the ROC space, the less accurate the test. **c**. A magnified view of the relationships between two predictor pairs. Hashlines demarcate the origin, data associated with iSPNs are uniquely situated on the lower-left corner of the graph (see inset). **d**. Left, A summary plot showing the mean and its 95% confidence interval for each of 23 variables for each of the two clusters. All variables were scaled. Right, t-SNE visualization of clustering. Cluster 1 (light gray) has a size of 26, whereas cluster 2 (dark gray) has a size of 30. Memberships for cluster 1: PV^+^ pre = 41.7% (5/12); PV^+^ light = 75.0% (9/12); Npas1^+^ pre = 56.3% (9/16); Npas1^+^ light = 18.8% (3/16). Memberships for cluster 2: PV^+^ pre = 58.3% (7/12); PV^+^ light = 25.0% (3/12); Npas1^+^ pre = 43.8% (7/16), Npas1^+^ light = 81.3% (13/16).

As noted earlier, as PV^+^ neurons and Npas1^+^ neurons produced diametric behavioral effects, we asked whether PV^+^ neurons and Npas1^+^ neurons could be working in concert to regulate the transitions between behavioral states in mice. Instead of simply providing granularity of behavioral dynamics, we sought to determine if we can use the different movement metrics collectively to define behavioral states. To this end, we parsed out the data from both PV-Cre and Npas1-Cre mice during both pre- and light-periods and then performed a *k*-means clustering (with *k* = 2) that included all 25 movement metrics. In **Figure 7d**, the dimensionality of the data is reduced via t-SNE, with each mouse shaded according to which cluster it is assigned to. As expected, the predicted cluster memberships for cluster 1 and 2 were about evenly split in both PV-Cre and Npas1-Cre mice during the pre-period (PV: 56.3%, Npas1: 41.7%). However, optogenetic stimulation shifted the membership distributions during the light-period—PV-Cre mice were predominantly (75.0%) in cluster 1, whereas Npas1-Cre were predominantly (81.3%) in cluster 2.

The *k*-means clustering analysis thus provided critical insights. First, it reinforced the idea that optogenetic manipulations of PV^+^ neurons and Npas1^+^ neurons give rise to distinct physiological behaviors. Second, the analysis suggests that the combinations of movement metrics can be used to define two behavioral states and supports the notion that PV^+^ neurons and Npas1^+^ neurons act as opposing forces that tune mouse behavior.

### Local collateral connectivity within the GPe is cell-specific

Based on the opposing roles of PV^+^ neurons and Npas1^+^ neurons, we hypothesized that they communicate with each other via inhibitory connections. Consistent with prior observations from single-cell labeling studies (Kita and Kitai, 1994; Nambu and Llinas, 1997; Bevan et al., 1998; Sato et al., 2000; Sadek et al., 2007; Fujiyama et al., 2016), biocytin-filled GPe neurons produced local collaterals that contain large varicosities that can be readily observed *post hoc* with confocal microscopy (**Figure 8a**). Although, the postsynaptic partner cannot be easily identified with this approach.

**Figure 8.**
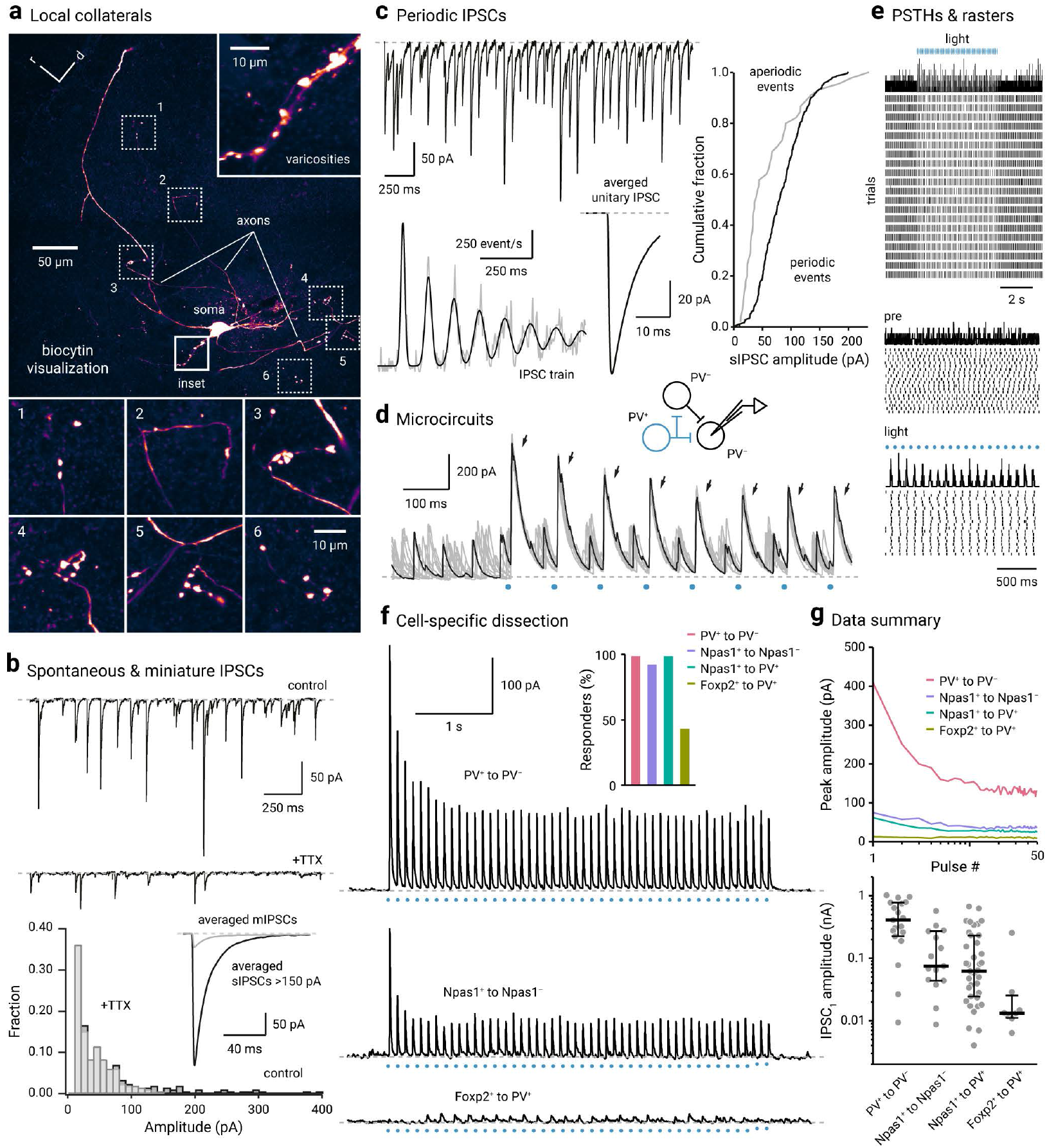
Local collateral connectivity within the GPe is cell-specific. **a**. An axon arborized locally in the vicinity of a biocytin-filled GPe neuron. Axons were identified as small-caliber processes that bear varicosities and contain no spines. Top right, an inset showing a series of varicosities produced by the biocytin-filled neuron. Bottom, Six additional examples of varicosities and their parent axons are shown. **b**. Top, Inwardly-directed spontaneous inhibitory postsynaptic currents (IPSCs) were readily observed in voltage-clamped GPe neurons *ex vivo*. Large, periodic events were blocked by the application of tetrodotoxin (1 µM), a voltage-gated sodium channel blocker. Bottom, A histogram showing the IPSC amplitude distribution. Inset, Averaged miniature IPSCs (gray) and large (> 150 pA) IPSCs (black) are shown. **c**. Top, A current trace showing periodic IPSCs recorded from a voltage-clamped GPe neuron. Bottom, An autocorrelation of the IPSC train from an example neuron with one unitary input, fitted with a sum of Gaussian components, indicating that the neuron received one periodic input in addition to aperiodic inputs (left). The waveform of the averaged unitary IPSC is shown (right). Right, A cumulative plot showing the difference in the amplitude distribution of periodic (black) and aperiodic events (gray). **d**. Spontaneous IPSCs and optogenetically evoked IPSCs (arrows) were observed from a voltage-clamped PV^−^ neuron (V_hold_ = –30 mV). A low-chloride internal solution was used. Light stimuli (blue circle) were delivered asynchronously relative to the occurrence of spontaneous IPSCs. A proposed microcircuit is shown on the top. **e**. Top, Peri-stimulus time histogram (PSTH) and raster plot showing the occurrence of spontaneous IPSCs. Evoked IPSCs were excluded. Middle & bottom, An expanded time scale showing the timing of spontaneous IPSCs was reset by evoked IPSCs. Periodicity is demonstrated by the evenly spaced peaks in the PSTH. No temporal structures in the spontaneous IPSCs were observed before light delivery. **f**. Local connectivity was dissected using optogenetics *ex vivo*. Example traces from PV^+^ to PV^−^ (top), Npas1^+^ to Npas1^−^ (middle), and Foxp2^+^ to PV^+^ (bottom) are shown. ChR2-eYFP negative neurons were targeted for recording. For positive identification of PV^+^ neurons, PV^+^ neurons were labeled with the PV-tdTomato allele. Inset, Four connection types were examined: PV^+^ to PV^−^ (blush, n = 19), Npas1^+^ to Npas1^−^ (purple, n = 16), Npas1^+^ to PV^+^ (green, n = 39), and Foxp2^+^ to PV^+^ (olive, n = 18). The proportion of neurons that responded are shown as bar graphs. Only a subset of recorded PV^+^ neurons (8 out of 18, 44.4%) showed detectable Foxp2^+^ input. **g**. Top, Population data showing IPSC amplitude for four different connection types across 50 pulses: PV^+^ to PV^−^ (blush, n = 19), Npas1^+^ to Npas1^−^ (purple, n = 15), Npas1^+^ to PV^+^ (green, n = 39), and Foxp2^+^ to PV^+^ (olive, n = 8). Each line represents median values. Bottom, Univariate scatter plot showing the first IPSC amplitudes for four different connection types. Thick lines represent the medians. Whiskers represent interquartile ranges.

The presence of local inhibitory networks was further supported by the presence of large, periodic inhibitory postsynaptic currents in voltage-clamp recordings with a high-chloride internal solution. As expected from the local rather than extrinsic (e.g., from the striatum) origin, these events are action potential-dependent, as demonstrated by their sensitivity to the application of tetrodotoxin (1 µM), a voltage-gated sodium channel blocker (**Figure 8b**). To gain further insights into the local connectivity, spontaneous IPSCs were recorded from PV^+^ neurons and Npas1^+^ neurons; properties of periodic and aperiodic IPSCs were extracted using our established analysis routine (Higgs and Wilson, 2016) (**Figure 8c**). In agreement with the results in **Figure 8b**, the small amplitude aperiodic events were likely of striatal origin, whereas the large amplitude periodic events likely arose from local collateral inputs. Furthermore, this analysis revealed that PV^+^ neurons and Npas1^+^ neurons receive local inputs and display IPSCs with very similar biophysical properties (**Table 5**).

**Table 5.** Properties of spontaneous IPSCs in PV^+^ neurons and Npas1^+^ neurons.

|  |  | PV<sup>+</sup> neurons |  | Npas1<sup>+</sup> neurons |  | Mann-Whitney<sup>b</sup> |
| --- | --- | --- | --- | --- | --- | --- |
|  |  | Median ± MAD | n<sup>a</sup> | Median ± MAD | n |  |
| All spontaneous inhibitory postsynaptic currents<sup>c</sup> | Rate (Hz) | 12.89 ± 3.63 | 50 | 7.76 ± 2.90 | 47 | 0.025 |
|  | Amplitude (pA) | 71.00 ± 20.54 | 50 | 60.24 ± 13.11 | 47 | 0.037 |
| Autocorrelation fit parameters | Aperiodic rate (Hz) | 8.01 ± 3.13 | 12 | 7.37 ± 1.34 | 10 | > 0.99 |
|  | Presynaptic frequency (Hz) | 8.88 ± 2.03 | 13 | 9.75 ± 1.14 | 11 | 0.49 |
|  | Periodic rate (Hz) | 6.69 ± 1.17 | 13 | 7.25 ± 1.50 | 11 | 0.56 |
|  | Presynaptic CV<sub>ISI</sub> | 0.13 ± 0.04 | 13 | 0.15 ± 0.06 | 11 | 0.28 |
|  | Power parameter | 0.70 ± 0.23 | 13 | 0.72 ± 0.10 | 11 | 0.88 |
| Unitary inhibitory postsynaptic currents | Amplitude (pA) | 133.64 ± 71.95 | 12 | 82.70 ± 40.25 | 10 | 0.20 |
|  | CV | 0.50 ± 0.16 | 12 | 0.56 ± 0.10 | 10 | 0.54 |
|  | Amplitude with failures (pA) | 111.91 ± 68.80 | 12 | 72.86 ± 39.60 | 10 | 0.25 |
|  | CV with failures | 0.80 ± 0.34 | 12 | 0.85 ± 0.11 | 10 | 0.82 |
|  | Variance/mean with failures | 79.00 ± 11.15 | 12 | 34.36 ± 10.50 | 10 | 0.15 |

<sup>a</sup>n equal to the number of neurons. <sup>b</sup>Two-tailed exact P values are listed. Abbreviations: CV = coefficient of variation, ISI = interspike interval, MAD = median absolute deviation, IPSCs = inhibitory postsynaptic currents. No results are significant after applying the Bonferroni correction at a significance level of 0.05.

While it is probable that PV^+^ neurons and Npas1^+^ neurons are reciprocally connected to each other via inhibitory connections, the precise local connectivity patterns between GPe neurons have not been examined previously. As termination sites produced by local axonal arborizations are sparse and spatially distributed within the GPe (Kita and Kitai, 1994; Nambu and Llinas, 1997; Bevan et al., 1998; Sato et al., 2000; Sadek et al., 2007; Bugaysen et al., 2013; Fujiyama et al., 2016), an optogenetic approach instead of paired recordings was used to more efficiently survey the local connectivity patterns. ChR2-eYFP was transduced into genetically-defined neurons in PV-Cre and Npas1-Cre mice, using a Cre-inducible approach. Transgene negative neurons were targeted for voltage-clamp recordings. As illustrated in **Figure 8d**, optogenetic stimulation of PV^+^ neurons induced antiphase timing of a spontaneous synaptic input, suggesting antiphase spiking of a putative PV^−^ neuron. This is more clearly shown in the peristimulus time histogram in **Figure 8e**. These results highlight the potential of local collateral input to shape population activity across the GPe. Using similar approaches, we tested the connectivity between various neuron types (PV^+^ to PV^−^, Npas1^+^ to Npas1^−^, Npas1^+^ to PV^+^, and Foxp2^+^ to PV^+^) (**Figure 8f** & **g**). To avoid confounds associated with photocurrents, ChR2-eYFP negative neurons were targeted for recording. We found that PV^+^ to PV^−^ input was the strongest of all connection types examined (PV^+^ to PV^−^: 409.9 ± 233.0 pA, *n* = 19 neurons; Npas1^+^ to Npas1^−^: 75.1 ± 59.2 pA, *n* = 15 neurons; Npas1^+^ to PV^+^: 62.2 ± 44.0 pA, *n* = 39 neurons; Foxp2^+^ to PV^+^: 13.3 ± 2.0 pA, *n* = 8 neurons; PV^+^ to PV^−^ vs. Npas1^+^ to Npas1^−^, P = 0.0024; PV^+^ to PV^−^ vs. Npas1^+^ to PV^+^, P < 0.0001; PV^+^ to PV^−^ vs. Foxp2^+^ to PV^+^, P = 0.00023; Npas1^+^ to Npas1^−^ vs. Npas1^+^ to PV^+^, P = 0.53; Npas1^+^ to Npas1^−^ vs. Foxp2^+^ to PV^+^, P = 0.0053; Npas1^+^ to PV^+^ vs. Foxp2^+^ to PV^+^, P = 0.0047) (**Table 6**). This finding is consistent with the earlier observation that PV^+^ boutons are abundant and form basket-like aggregates on cell bodies of both PV^+^ and PV^−^ neurons within the GPe (Kita, 1994). Though Npas1^+^ neurons did target PV^+^ (which are Npas1^−^) neurons, the strength was smaller than that of the PV^+^ to PV^−^ connection. The asymmetry in the connectivity between PV^+^ neurons and Npas1^+^ neurons can be partially explained by their abundance. Additional factors, such as the number of release sites, quantal content, and postsynaptic receptor density may also be at play.

**Table 6.** Local collateral connectivity within the GPe.

| Input | IPSC measure |  |  | Statistical analyses<sup>b</sup> |  |  |  |
| --- | --- | --- | --- | --- | --- | --- | --- |
| | Median ± MAD (pA) | Response rate (%) | n<sup>a</sup> | vs. PV<sup>+</sup> to PV<sup>-</sup> | vs. Npas1<sup>+</sup> to Npas1<sup>-</sup> | vs. Npas1<sup>+</sup> to PV<sup>+</sup> | vs. Foxp2<sup>+</sup> to PV<sup>+</sup> |
| PV<sup>+</sup> to PV<sup>-</sup> | 409.9 ± 233.0 | 100 | 19 | - | **0.0024** | **< 0.0001** | **0.00023** |
| Npas1<sup>+</sup> to Npas1<sup>-</sup> | 75.1 ± 59.2 | 93.8 | 16 | **0.0024** | - | 0.53 | **0.0053** |
| Npas1<sup>+</sup> to PV<sup>+</sup> | 62.2 ± 44.0 | 100 | 39 | **< 0.0001** | 0.53 | - | **0.0047** |
| Foxp2<sup>+</sup> to PV<sup>+</sup> | 13.3 ± 2.0 | 44.4 | 18 | **0.00023** | **0.0053** | **0.0047** | - |

<sup>a</sup>Sample size (n) included all cells with or without responses. <sup>b</sup>Mann-Whitney U test on peak amplitude; two-tailed exact P values were shown. The bolded P values are significant when using a Bonferroni correction at a significance level of 0.05, adjusting for the fact that 6 comparisons were considered.

Only a subset of recorded PV^+^ neurons (8 out of 18, 44.4%) showed detectable Foxp2^+^ input, and the strength of this connection was the weakest among all connection types examined (Foxp2^+^ to PV^+^ vs. PV^+^ to PV^−^, P = 0.00023; Foxp2^+^ to PV^+^ vs. Npas1^+^ to Npas1^−^, P = 0.0053; Foxp2^+^ to PV^+^ vs. Npas1^+^ to PV^+^, P = 0.0047) (**Table 6**). The observation of this weak and likely biologically negligible connection is highly consistent with recent studies (Aristieta et al., 2020; Ketzef and Silberberg, 2020). As it was clear that the Npas1^+^ input targets PV^+^ neurons, these results collectively imply that PV^+^ neurons receive Npas1^+^ input largely from the Nkx2.1^+^ (aka Npr3^+^) subpopulation, which do not express Foxp2 (**Figure 1c**), see also (Abecassis et al., 2020).

## Discussion

In this study, we determined that Npr3^+^ neurons shared the unique molecular and anatomical signature of Npas1^+^-Nkx2.1^+^ neurons. While Kcng4^+^ neurons share canonical properties with PV^+^ neurons, they were not involved in regulating anxiety-like behavior (as measured with center-edge preferences). By using machine learning and statistical approaches to delineate the functional properties of GPe neuron subtypes, we proposed that PV^+^ neurons and Npas1^+^ neurons work in concert to control the transitions between behavioral states in mice. This idea was further supported by the existence of local connectivity between them.

### GPe neuron diversity

What is a cell type, and why is it an important question? Both the embryonic origins and the congruent expression of transcription factors govern cell specifications (Hu et al., 2017; Lim et al., 2018; Fishell and Kepecs, 2020). It is well-established that the GPe primarily arises from the medial and lateral ganglionic eminence (MGE and LGE) (Flandin et al., 2010; Nobrega-Pereira et al., 2010; Dodson et al., 2015; Chen et al., 2017). It is clear that Npas1^+^ neurons are derived from both progenitor pools; Foxp2^+^ neurons are from the LGE, whereas Npr3^+^ neurons are from the MGE and maintain the expression of Nkx2.1 (**Figure 1c**). Consistent with the single-cell transcriptomic data (Saunders et al., 2018), our current and past (Abecassis et al., 2020) data showed the molecular complexity of PV^+^ neurons (**Figure 9**). This is in agreement with earlier studies that hinted at multiple MGE progenitors giving rise to at least two populations of PV^+^ neurons (Silberberg et al., 2016). In fact, heterogeneity among PV^+^ neurons can be a general feature across brain areas (Dehorter et al., 2015; Garas et al., 2016; Tasic et al., 2018; Hodge et al., 2019; Huang and Paul, 2019; Mahadevan et al., 2020). It has been shown that there is sonic hedgehog (Shh) expression in the preoptic area and in the nascent GPe (Flandin et al., 2010; Sousa and Fishell, 2010). In agreement with this, earlier studies on Dbx1^+^ neurons showed that preoptic area neurons make a small contribution to the cellular makeup of the GPe (Gelman et al., 2011; Abecassis et al., 2020). This inference was confirmed with our recent, progenitor-specific fate-map study (Poulin et al., 2020).

**Figure 9.**
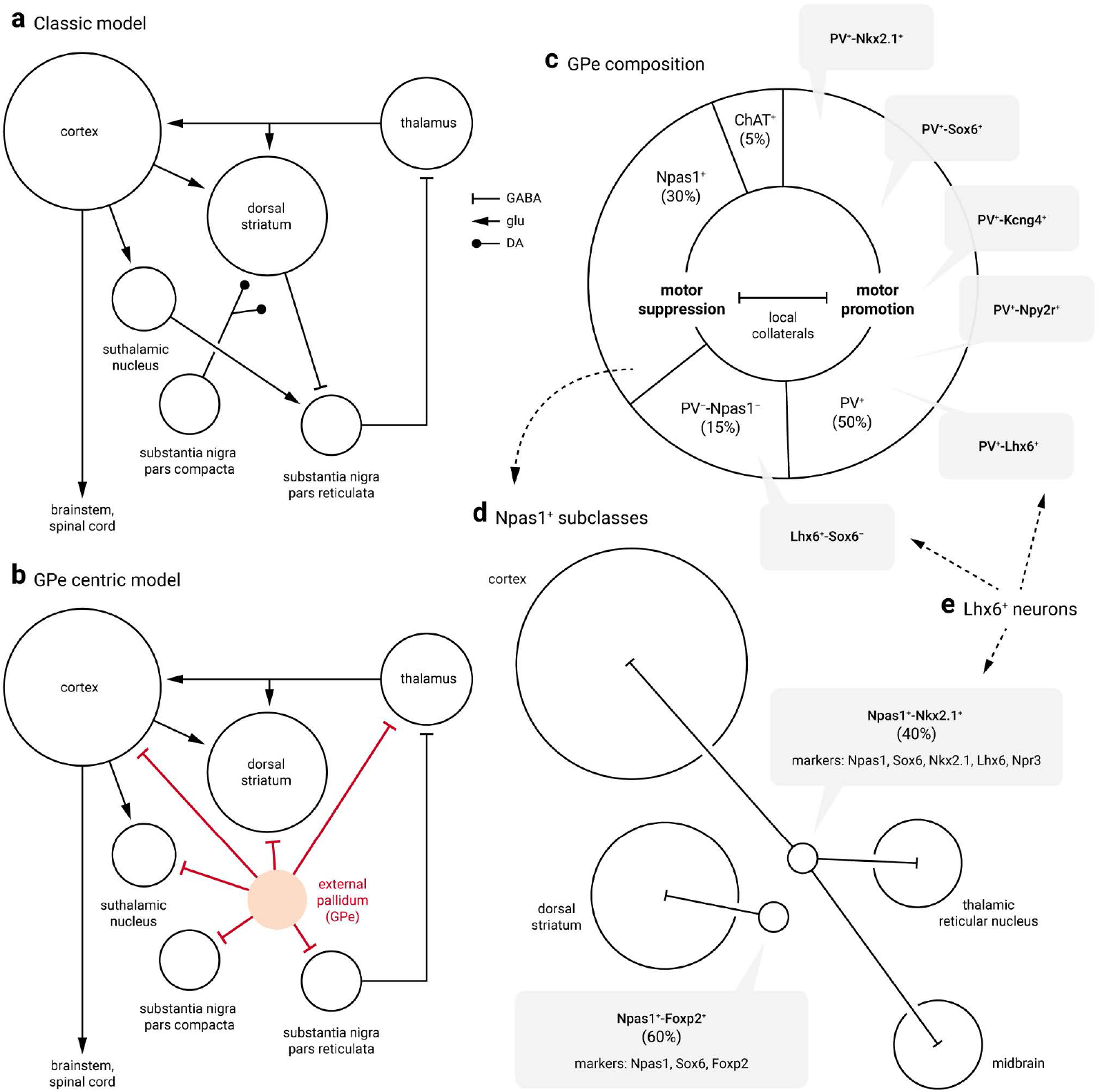
Cellular and circuit substrates for motor regulation by the GPe. **a**. A simplified circuit diagram showing the key nodes within the basal ganglia. The dStr and STN are the two input stations that receive excitatory cortical inputs; the SNr is the output station that provides inhibitory projection to the thalamus. The entire basal ganglia circuit, especially the dStr, is under the neuromodulatory control of the SNc. Size of the target areas (circles) is an artistic rendering based on the volume of the brain areas. The circuit role of the GPe is underappreciated. The GPe was traditionally viewed as a “black box,” which relays information from the dStr to the STN. **b**. A modified circuit model highlighting the central role of the GPe within the basal ganglia. The GPe sends inhibitory projections to all key structures within the circuit. **c**. Pie chart summarizing the neuronal composition of the mouse GPe. The area of the sectors represents the approximate size of each neuron class. PV^+^ neurons (which constitute 50% of the GPe) are heterogeneous. Nkx2.1, Sox6, Kcng4, Npy2r, and Lhx6 are coexpressed in PV^+^ neurons to a varying extent. How they intersect with each other remains to be determined. Previous transcriptomic data point to the idea that Kcng4^+^ neurons represent a unique subclass of PV^+^ neurons in the GPe. By projecting to the STN and SNr, PV^+^ neurons assume the canonical role of the GPe. Npas1^+^ neurons are 30% of the GPe; they can be subdivided into two subclasses (see **d**). Consistent with the distinct electrophysiological properties, PV^+^ neurons and Npas1^+^ neurons have unique behavioral roles: PV^+^ neurons are motor-promoting, while Npas1^+^ neurons are motor-inhibiting. PV^+^ neuron and Npas1^+^ neuron classes jointly tune motor behavior through a push-pull mechanism. Local inhibitory connections between these neuron classes serve as key cellular substrates that underlie this process. ChAT^+^ neurons are ∼5% of the total GPe neuron population and show no overlap with other known classes of GPe neurons. They project broadly to cortical areas. PV^-^-Npas1^-^ neurons amount to ∼15% of the total GPe and are the least characterized neuron subtype (see **e**). **d**. Two bona fide subclasses of Npas1^+^ neurons are identified in the mouse GPe. They differ in their molecular marker expression, axonal projections, and electrophysiological properties. Npas1^+^-Foxp2^+^ neurons represent 60% of the total Npas1^+^ neuron class; they project exclusively to the dStr, targeting SPNs. Npas1^+^-Npr3^+^ (aka Npas1^+^-Nkx2.1^+^ and Npas1^+^-Lhx6^+^) neurons represent 40% of the total Npas1^+^ neuron class; they project to the cortex, thalamus, and midbrain. Npas1^+^-Foxp2^+^ neurons alone are sufficient to account for the motor-suppressing effect produced by optogenetic stimulation of pan-Npas1^+^ neurons *in vivo*. In contrast, the behavioral roles of Npas1^+^-Npr3^+^ awaits further clarification. Size of the target areas (circles) is an artistic rendering based on the volume of the target areas, and does not reflect the axonal density, synaptic strength, or contacts formed by Npas1^+^ neuron subclasses. **e**. Lhx6^+^ neurons are heterogeneous. They encompass three different subpopulations of neurons, including those in the PV^+^ and Npas1^+^ neuron classes. Both PV^+^-Lhx6^+^ neurons and Npas1^+^-Npr3^+^ neurons coexpress Sox6. Lhx6^+^-Sox6^-^ neurons are a subset of PV^-^-Npas1^-^ neurons.

Although cellular diversity in the GPe has been recognized since the 1970’s (DeLong, 1971; Fox et al., 1974), how heterogeneity maps onto molecular and functional differences has been unclear. Our interest was never to understand cell-type *per se*. Instead, by studying cellular composition and the nature of neuronal identity, we hope to uncover molecular markers and genetic tools that allow us to interrogate the organization principles and functions of GPe neuron subtypes. The cellular composition in the GPe is relatively simple compared to that of the cortex and hippocampus (Rudy et al., 2011; Jiang et al., 2015; Tasic et al., 2016; Tasic et al., 2018; Huang and Paul, 2019; Adkins et al., 2020; Gouwens et al., 2020; Yao et al., 2020); the advent of single-cell transcriptomic and mouse genetics has created an opportunity to interrogate the GPe with previously unexplored tools and has enabled us to provide new insights into GPe neuron diversity and its behavioral relevance in this study (**Figure 9**). A large set of transcription factors have been previously identified to govern the formation of the GPe. We took a leap of faith in generating Npas1-Cre mice and Npas1 antibodies *de novo* to interrogate the GPe. This decision was based on the compelling data from prior literature (Flandin et al., 2010; Nobrega-Pereira et al., 2010). While PV-Cre and Npas1-Cre mice will continue to be instrumental to our research, we have now identified a set of transgenic lines that will allow the field to work together to solve more challenging questions and to reconcile discrepancies that arose from suboptimal tools.

Depending upon the metric employed, the definition of cell types differs. This is our first attempt to examine the cellular, circuit, and behavioral properties of Foxp2^+^ neurons, Npr3^+^ neurons, Kcng4^+^ neurons, and Npy2r^+^ neurons. This study is among the first to show that GPe neuron subclasses have specialized connections and functions (**Figure 9**). However, we do not know the relationship between these neuron subtypes and the somatotopic organization of the circuit (Romanelli et al., 2005; Nambu, 2011; Bertino et al., 2020). *In vivo* optogenetics does not offer the resolution to answer this type of question. Our understanding of these neurons, and ultimately the GPe, will undoubtedly continue to be refined, while we await driver lines that offer genetic access to the PV^+^-Kcng4^−^ population.

### Beyond go and no-go?

By tracking body kinematics and movement dynamics, we described how optogenetic manipulations of specific cell types gave rise to unique behavioral structures. With this information, we inferred the behavioral states of mice and functional interactions between individual neuron types. We found that upon Npas1^+^ neuron stimulation, mice were not completely motionless. Instead, they spent more time executing fine movement, which on its own can be mediated by a complex circuit. As motionless and fine movement often co-occurred within a given time period, we interpret fine movement as a manifestation of action suppression. In turn, the basal ganglia are reset for selecting new actions—in this case, locomotion in an open-field arena.

As PV^+^ neurons and dSPNs converge onto the substantia nigra pars reticulata (Grofova, 1975; Araki et al., 1985; Bolam et al., 1993; Mink, 1996; Smith et al., 1998; Connelly et al., 2010), it is conceivable that PV^+^ neurons and dSPNs are both involved in motor-promotion. Anatomical studies show that SNr-projecting striatal neurons arborize within the GPe (Kawaguchi et al., 1990; Wu et al., 2000; Levesque and Parent, 2005; Fujiyama et al., 2011; Cui et al., 2020); however, the postsynaptic targets remain elusive. Our data suggest that they do not target PV^+^ neurons. Though the role of the basal ganglia in movement speed has long been recognized (Kornhuber, 1971; DeLong et al., 1984; Yttri and Dudman, 2016; Fobbs et al., 2020), here we argue that the basal ganglia control more than just go and no-go. As the basal ganglia are known to support a wide repertoire of innate complex behaviors (such as rearing and grooming) (Graybiel, 2008; Redgrave et al., 2010; Ahmari, 2016; Markowitz et al., 2018; Park et al., 2020), we should consider using movement dynamics broadly as metrics to measure basal ganglia function rather than relying solely on speed. Our experiments utilized different mouse strains to target selective neuron types for manipulations, and therefore, it is rather difficult to untangle the *bona fide* cell-specific behavioral effects from nuanced differences in behavior across mouse strains. Future experiments that employ intersectional approaches will yield more definitive answers.

We have recently shown that PV^+^ neurons and Npas1^+^ neurons are positioned differently within the basal ganglia. This inference is further supported by *in vivo* whole-cell analysis (Ketzef and Silberberg, 2020). We did not observe marked differences in the electrophysiological properties between neurons within each class. This implies that input-output transformation among them is similar. It is thus probable that neurons within each class receive inputs with similar physiological properties. In turn, both PV^+^ neurons and Npas1^+^ neurons exhibit complex projection patterns. It is now clear that this complexity is in part a result of a mixture of neuron subclasses (Mastro et al., 2014; Abdi et al., 2015; Dodson et al., 2015; Hernandez et al., 2015; Fujiyama et al., 2016; Glajch et al., 2016; Hegeman et al., 2016; Abecassis et al., 2020). The formation of these “circuit modules” by neuron subclasses, in turn, dictates the network effects and thus behavioral outcome. We previously examined the synaptic interactions between GPe neurons and key structures within the basal ganglia—the dorsal striatum and the STN (Chan et al., 2004; Fan et al., 2012; Hernandez et al., 2015; Cui et al., 2016; Glajch et al., 2016; Cui et al., 2020; Pamukcu et al., 2020). Given the diametric behavioral effects produced by these two populations of neurons, it was a logical next step to ask if PV^+^ neuron and Npas1^+^ neurons are reciprocally inhibiting each other. In this study, by using a combination of transgenic lines, we have overcome some of the challenges associated with the interrogation of cell type-specific connectivity. Our data showed that PV^+^ neurons and Npas1^+^ neurons are connected to each other and thus support the idea that the diametric behavioral phenotype observed when one of these two classes of neurons was perturbed was a result of a push-pull function between PV^+^ neurons and Npas1^+^ neurons. We found that Foxp2^+^ neurons completely avoided PV^+^ neurons. More importantly, this observation suggests that input from Npas1^+^ neurons to PV^+^ neurons are provided by Npr3^+^ (aka Npas1^+^-Nkx2.1^+^) neurons exclusively. This inference is consistent with the high density of local axons produced by Npr3^+^ neurons, as observed in **Figure 1c**. However, we have yet to uncover the properties of the connections made by Npr3^+^ neurons. In contrast, synaptic tagging experiments suggested that Kcng4^+^ neurons did not produce local inputs (**Figure 1c**). It will be important to confirm this anatomical observation with functional studies. With the identification of new genetic tools, we should be in a position to study the synaptic connectivity and its alterations at greater depth.

Recent literature emphasized that PV^+^ neurons are at the intersection of motor-anxiety circuits (Sztainberg et al., 2011; Hunt et al., 2018; Giovanniello et al., 2020); we do not currently know the downstream targets that mediate the anxiety-related response. As the STN does not mediate this anxiety-related response, other descending pathways such as the direct projections to the raphe (Pollak Dorocic et al., 2014) or their indirect projection to the habenula via the internal globus pallidus (Smith et al., 1998; Wallace et al., 2017) could be involved. While Npas1^+^ neurons, in particular Foxp2^+^ neurons, have extensive axonal arborization in the dorsal striatum (Dodson et al., 2015; Hernandez et al., 2015; Glajch et al., 2016), it is important to emphasize that their downstream impact on the somatic excitability of striatal projection neurons is relatively weak (Glajch et al., 2016). The cellular mechanisms that mediate motor-suppression have yet to be identified. While Foxp2^+^ neurons have been proposed to send “stop” signals to the striatum (Mallet et al., 2016; Aristieta et al., 2020; Goenner et al., 2020), it is likely that multiple circuit elements are involved in producing complete behavioral arrest (Schmidt et al., 2013; Jahanshahi et al., 2015; Wessel and Aron, 2017). On the other hand, it is clear that the inhibitory projection to the STN (and substantial pars reticulata) mediates the motor-promoting effects, and it is consistent with the established relationship between STN activity and movement suppression (Hamani et al., 2004; Aron and Poldrack, 2006; Aron et al., 2007; Eagle et al., 2008; Schmidt et al., 2013; Schweizer et al., 2014; Fife et al., 2017; Wessel and Aron, 2017). In addition to the established literature demonstrating that the GPe sends inhibitory signals to the input and output nuclei of the basal ganglia (Jessell et al., 1978; Mink, 1996; Smith et al., 1998; Kita, 2007; Hegeman et al., 2016; Adam et al., 2020), recent studies highlight the existence of cortico-pallido-cortical loops (Naito and Kita, 1994; Chen et al., 2015; Saunders et al., 2015; Schwarz et al., 2015; Ahrlund-Richter et al., 2019; Karube et al., 2019; Abecassis et al., 2020; Adkins et al., 2020; Anastasiades et al., 2020; Clayton et al., 2020; Gehrlach et al., 2020; Lee et al., 2020; Muñoz-Castañeda et al., 2020; Garcia et al., 2021). However, the precise circuitry has yet to be defined. The direct feedback from Npr3^+^ (aka Npas1^+^-Nkx2.1^+^) neurons to cortical regions (**Figure 1c**, see also (Abecassis et al., 2020)) argues that action selection is not simply a top-down command.

## Acknowledgments

We thank Daniela Garcia, Alyssa Bebenek, Moises Melesio, Kris Shah, Ahana Narayanan, Vaishnavi Tetali, Saivasudha Chalasani, and Daniel Hegeman for their assistance on the project, Dr. Tiffany Schmidt for providing Kcng4-Cre mice, Dr. Byung Kook Lim for providing the synaptophysin-GFP virus, and Drs. Ann Kennedy and Sam Golden for input on the behavioral analyses. This work is supported by NIH R01 NS069777 (CSC), R01 MH112768 (CSC), R01 NS097901 (CSC), R01 MH109466 (CSC), R01 NS088528 (CSC), T32 AG020506 (AP), T32 NS041234 (HSX), F32 NS098793 (HSX), R01 MH112768 (NJJ), R35 NS097185 (CJW), HHMI-PF Medical Research Fellowship (ZAA), and AΩA Student Research Fellowship (ZAA).

## Author contributions

SC and BLB performed histological analysis. SC, QC, IYMC, HSX, YZ, HM, ZAA, and XD surveyed the intrinsic properties. QC, HMH, XD, YZ, and CJW examined the local connectivity. AP, QC, SR, and NJJ conducted the behavioral studies. IYMC developed the machine learning analysis pipeline. YC and SMB provided expertise for neuron classification. CSC wrote the manuscript with input from all co-authors. All authors reviewed and edited the manuscript. CSC designed, directed, and supervised the project.

## Video Legends

Supplemental video. Machine learning approaches capture mouse behavior motifs in an open field. Representative movie of a mouse in an open field arena. Current behavior of the mouse is displayed. Event timers track the cumulative duration of locomotion, rearing, fine movement, and motionless events. Video is shown at 0.5x speed.

## Notes

https://graceyichen.shinyapps.io/neuro_interactive_analysis/

### Competing Interest Statement

The authors have declared no competing interest.

